# Developing potent and selective TBK1 molecular glue degraders for cancer immunotherapy

**DOI:** 10.64898/2026.01.30.702304

**Authors:** John J. Caldwell, Patrick R. A. Zanon, Iona Ferguson, Stephen T. Hallett, Tencho Tenev, Natalia Serrano-Aparicio, Russell J. H. Wood, Shakil Khan, Sidonie Wicky John, Rebecca Wilson, Matthew T. Warren, P. Craig McAndrew, Martin Steger, Denise Winkler, Uli Ohmayer, Bjoern Schwalb, Bachuki Shashikadze, Yann-Vaï Le Bihan, Andrea Scarpino, Konstantinos Mitsopoulos, Fiona Dziegiel, Agnieszka Konopacka, Rob L. M. van Montfort, Pascal Meier, Henrik Daub, Zoran Rankovic

## Abstract

Immune checkpoint blockade (ICB) has transformed cancer therapy across multiple tumour types, yet primary and acquired resistance remain major barriers to durable benefit. TANK-binding kinase 1 (TBK1) has emerged as an attractive target to enhance anti-tumour immunity, acting as a serine/threonine kinase that restrains immunogenic cell death and downstream immune activation. Here we report the discovery of a first-in-class TBK1 molecular glue degrader (MGD), CCT412020, identified through high-throughput proteomics screening of a next-generation molecular glue library. CCT412020 induces rapid, potent and selective TBK1 degradation across a panel of breast cancer cell lines. A cryo-EM structure of CCT412020 in complex with CRBN/ΔBPB-DDB1 and TBK1-homodimer reveals an unexpected binding mode that bypasses the canonical G-loop and instead engages an unconventional site at the TBK1 homodimer interface. Functionally, TBK1 loss via CCT412020 sensitises tumour cells to TNF- and interferon-driven responses and reduced viability across a broad range of cancer cell lines. Together, these findings establish CCT412020 as a mechanistically distinct TBK1 degrader and provide a framework for developing TBK1-targeted degraders as immunomodulatory anti-cancer agents to overcome ICB resistance.

## Introduction

Immune checkpoint blockade (ICB) has transformed cancer therapy by releasing inhibitory pathways that restrain anti-tumour T cell activity, thereby enabling immune recognition and elimination of malignant cells^1^. However, while ICB is now the standard-of-care across multiple tumour types, many patients fail to respond or relapse, largely due to cancer cells developing evasive mechanisms^2^. Across cancers that are considered ICB-responsive, objective response rates typically range from 20% - 40%^3^, and there are no approved strategies that reliably overcome primary or acquired resistance once it emerges, underscoring a major unmet clinical need. To address this challenge and identify tumour-intrinsic drivers of resistance to immunotherapy, unbiased target-discovery approaches have been deployed to identify tumour-intrinsic regulators of immunotherapy response, most prominently loss-of-function CRISPR-Cas9 screening *in vitro* and *in vivo*^4, 5^. These efforts have converged on a limited set of actionable nodes that shape tumour-immune interactions. Among the most compelling is TANK-binding kinase 1 (TBK1)^6^, a serine/threonine kinase positioned at the interface of innate immune signalling and stress responses^7^, that is increasingly implicated as a tumour cell-intrinsic constraint on immunogenicity and productive anti-tumour immunity.

TBK1 is a member of the noncanonical inhibitor of nuclear factor-κB (IκB) kinase (IKK) family of serine/threonine kinases^8^, which also includes the closely related homologous IκB kinase subunit ε (IKKε). Both kinases play important regulatory roles in innate immune response to bacterial and viral challenges^9^. In this process, activated by pathogen-associated molecular patterns (PAMPs) and inflammatory cytokines such as tumour necrosis factor (TNF), TBK1 and IKKε directly phosphorylate and activate interferon regulatory factor 3 (IRF3) and IRF7 transcription factors. Phosphorylated IRFs form homo- and hetero-dimers and translocate into the nucleus where they induce interferon-stimulated genes (ISGs) that are critical to the host immune response.

In addition to its established functions in innate immune signalling, TBK1 has recently been shown to prevent TNF-induced cell death by phosphorylating and inhibiting receptor-interacting serine/threonine-protein kinase 1 (RIPK1), a key regulator of inflammation, cell survival and cell death^4, 10, 11, 12, 13, 14^. TNF stimulation can trigger two opposing outcomes: pro-survival inflammatory gene expression or cell death, driven by two RIPK1-containing signalling complexes^15, 16^. Upon TNF binding to TNF receptor 1 (TNFR1), a plasma membrane-associated complex-I forms, comprising adaptor proteins, E3 Ubiquitin ligases and kinases, including RIPK1. Ubiquitination of components of complex-I by cIAP1 and cIAP2 E3 ligases promotes recruitment of transforming growth factor-β-activated kinase 1 (TAK1) and the linear ubiquitin chain assembly complex (LUBAC), which extends linear ubiquitin chains that serve as docking sites for nuclear factor-κB essential modulator (NEMO). NEMO, together with TANK and NAK-associated protein 1 (NAP1), coordinates the recruitment of canonical IKKs (IKKα/IKKβ) as well as TBK1 and IKKε. Activation of TAK1 and the canonical IKKs within complex I drives NF-κB-dependent inflammatory gene expression. If complex I is destabilised, components including RIPK1 can relocalise to the cytosol to form complex II, which recruits caspase-8 via FADD to trigger apoptosis, or RIPK3 to initiate necroptosis. While TBK1 and IKKε are dispensable for transcriptional activation downstream of TNFR1, they are critical for restraining RIPK1 by preventing its autophosphorylation and subsequent transition into complex II^7, 10, 12, 17, 18^. RIPK1 autophosphorylation is thought to induce conformational changes that expose its death domain and RIP homotypic interaction motif, enabling engagement of FADD and RIPK3, respectively^16, 19, 20, 21, 22^. Thus, by blocking caspase-8- and RIPK3-dependent immunogenic cell death pathways, TBK1 functions as a molecular brake on immunogenic cell death and the downstream anti-tumour immune response.

Consistent with these findings, accumulating evidence over the past few years has implicated TBK1 in tumorigenesis^23^. Although recurrent TBK1 mutations are uncommon in human cancers, elevated TBK1 expression and/or dysregulated TBK1 activity has been reported across multiple malignancies, including in non-small cell lung cancer (NSCLC), pancreatic ductal adenocarcinoma (PDAC), cholangiocarcinoma, clear cell renal cell carcinoma (ccRCC), adult T-cell leukaemia, melanoma, oesophageal cancer, and breast cancer, amongst others^6, 23, 24, 25^. TBK1 may promote cancer initiation and progression through several non-mutually exclusive mechanisms, including supporting tumour cell survival and proliferation, as well as dampening anti-tumour immunity, for example by increasing immune checkpoint ligand expression and sustaining an immunosuppressive programme within the tumour microenvironment^23, 26^. TBK1 has emerged as a tumour cell-intrinsic immune evasion factor that functions as an intracellular “cell death checkpoint” in cancer cells ^7, 8, 10, 12, 27^. Loss of TBK1 primes tumour cells to undergo RIPK1- and caspase-dependent cell death in response to effector cytokines such as TNF and IFNγ. Importantly, TBK1 depletion does not appear to drive major remodelling of the immune compartment, but instead it lowers the threshold for tumour cell killing by TNF and IFNγ, thereby enhancing sensitivity to immune-mediated cytotoxity^23^. Consistently, genetic TBK1 loss in *in vitro* and *in vivo* cancer models effectively sensitises tumours to ICB treatment^7, 8, 10,12^. Taken together, these findings support TBK1 as a compelling therapeutic target to enhance anti-tumour immunity and overcome resistance to cancer immunotherapy.

Another groundbreaking development in drug discovery over the recent years is the emergence of targeted protein degradation (TPD) approaches, such as proteolysis targeting chimeras (PROTACs)^28, 29^ and molecular glue degraders (MGDs)^30^. MGDs are small molecules designed to hijack normal cellular ubiquitin-proteasome system (UPS) to degrade disease causing proteins. The immunomodulatory imide drugs (IMiDs) thalidomide, lenalidomide, and pomalidomide were the first drugs found to exert their pharmacological effect by inducing protein degradation. IMiDs were found to bind cereblon (CRBN), the substrate recognition receptor of the E3 Ubiquitin ligase complex CRL4^CRBN^, and recruit neosubstrates such as lymphoid transcription factors IKZF1 and IKZF3, whose ubiquitination and subsequent degradation underpins the clinical efficacy of these drugs in multiple myeloma^31, 32^. Since these initial findings, IMiDs were found to induce degradation of dozens of other neosubstrates^33^. Interestingly, despite their close structural similarity, IMiDs display different protein degradation profiles. For example, lenalidomide induces degradation of CK1α protein, while thalidomide does not^34^. Following this observation, chemical diversification approaches around the IMiD scaffolds that emerged in recent years proved to be a successful strategy for discovering potent and selective MGDs of novel neosubstrates with promising translational value^35, 36, 37^. Most CRBN neosubstrates were found to share a structural motif comprised of a β-hairpin loop featuring a conserved glycine residue at its apex, known as a G-loop^38, 39^ The G-loop functions as a recognition degron motif that interacts with a hotspot on the CRBN surface. Although several neosubstrates were reported to engage with CRBN in a different binding mode, the G-loop is generally considered as the CRBN canonical degron motif^40^. Computational mining of the human proteome identified more than 1,600 proteins harbouring G-loop-like motifs, further highlighting drug discovery opportunities^41^. Indeed, we and others have demonstrated that the CRBN neosubstrate landscape is much larger than the one defined by the first generation IMiDs, presenting great opportunities for translational research^40, 42, 43^. Encouraged by these promising initial findings we employed state-of-the-art proteomics technologies and the latest molecular glue library design strategies to develop our next generation MGD discovery platform. Here we report the discovery and extensive characterisation of CCT412020, the first-in-class CRBN-recruiting MGD with an unusual binding mode.

## Results

### Developing the next-generation Library of Molecular Glue Degraders

While large strides have been made towards *de novo* design of molecular glues, library screening remains essential. The governing design principles for first-generation molecular glue libraries (MGLs) were mostly based on maximising diversity around IMiD scaffolds^44^. This approach provided many successful outcomes and generated invaluable early understanding of the degron motif, MGD binding, and preliminary data on structure-degradation relationship (SDR). Fundamental to the design of our next generation molecular glue library (MGL.v2) was to fully leverage knowledge that emerged over the recent years, and move from a randomly applied chemical diversity to a more focused, knowledge-based exploration of molecular structure and property space (**Fig. 1A**). This “focused diversity” approach relies on in-depth analysis of literature reports and internal data to generate and incorporate knowledge-driven hypotheses into the library design. For example, some of the earlier literature reports suggest that exit vectors can dramatically influence degrader proteome-wide selectivity^45^. Analogues with C5 modifications on the phthalimide ring were found to display more selective profiles with reduced degradation of the C2H2 family of zinc finger (ZF)-containing proteins relative to analogues with identical modifications on the C4 position^45^. Computational modelling suggested that C5 modifications are more likely to create a steric clash with the ZF than C4 modifications.

**Figure 1.**
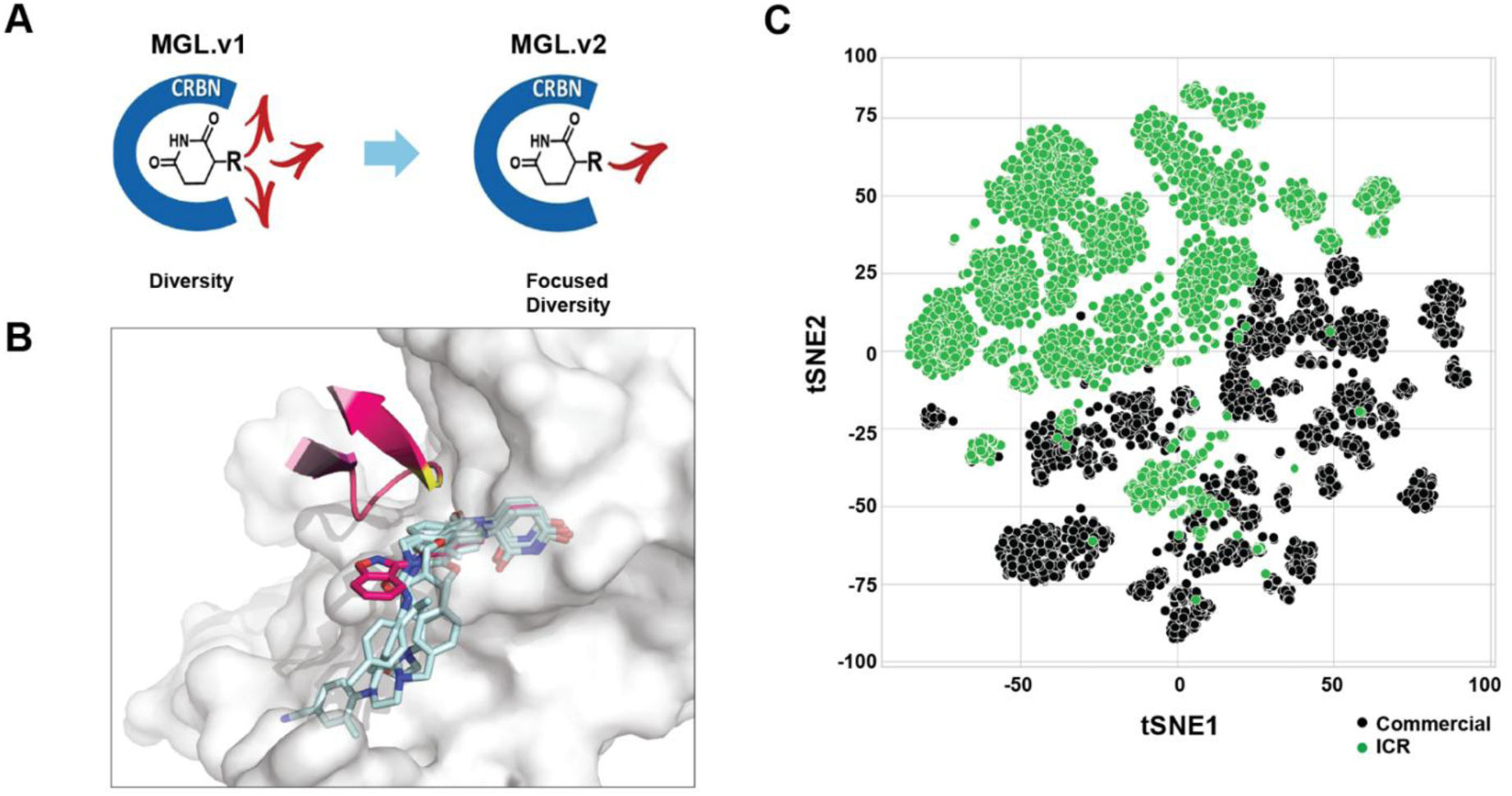
Molecular Glue Library (MGL.v2) design principles. **(A)** Schematic representation of library design hypothesis focused on diversity on a selected exit vector of the CRBN pocket space. **(B)** Alignment of the CK1α:SJ3149:CRBN ternary complex with a set of known β-hairpin-G-loop molecular glue degraders published along with their respective ternary complexes. Structures have been aligned on the residues of the CRBN binding site. The CK1α degron is depicted in magenta, with the glycine residue highlighted in yellow. CRBN molecular surface is represented in white. Molecular glue molecules presented are, SJ3149 in magenta and lenalidomide, CC-885, CC-90009, CC-220 (iberdomide) and CC-92480 (mezigdomide) in light blue (PDB IDs: 8G66, 5FQD, 5HXB, 6XK9, 8D80 and 8RQC, respectively). The alignment shows how well known MGDs interact along the CRBN surface, while SJ3149 exits the CRBN binding site towards the degron, interacting with the protein of interest. **(C)** t-SNE 2D decomposition of the library chemical space in comparison to the commercially available Enamine Ltd. IMiD library^49^.

Indeed, our inspection of ternary complex structures available in the Protein Data Bank (PDB)^46^ showed that various MGDs with C4 substituents form contacts mostly with the residues lining the CRBN surface, as exemplified by the structure of CC-220 in the complex with CRBN and IKZF1 (in light blue, **Fig. 1B**)^47^. With this in mind, we hypothesised that CRBN binders designed for a more extensive engagement with the neosubstrates’ degron motif may create a better chance of discovering potent and selective degraders of novel CRBN neosubstrates. Consequently, preference was given to CRBN binders with a C5-like vector that projects towards and along the degron G-loop. Indeed, this hypothesis inspired the “C4-to-C5-switch” strategy that we successfully used earlier to optimise a screening hit into a potent and selective CK1α degrader, SJ3149^48^ (in magenta, **Fig. 1B**). Based on this hypothesis, we designed a set of “degron-targeting” scaffolds and functionalised them in a single step using a diverse set of building blocks. One of the consequences of this degron-targeting design approach is that it leads to larger molecules. For that reason, we relaxed the cutoff for molecular size to 650 Da, while retaining the rest of the physicochemical properties within the traditional drug-like space.

Another hypothesis was based on our observation that phenyl glutarimide (PG) scaffold, which we initially developed as a CRBN warhead for PROTAC development^50^, often produce highly selective MGDs^42^. Hence, PG scaffolds and their variations were incorporated into the library. In addition, to filter out potential degraders of known neosubstrates such as GSPT1 and CK1α, we applied in our design process *in silico* models developed using our extensive internal structural and biochemical data. The final compounds were synthesised at >95% purity and plated in a 384-well format for screening. The library expansion has been a continuous and highly dynamic process. As new insights emerged from the screening campaigns, internal programs and literature, new hypotheses were created and incorporated into the library. Building on this knowledge-driven multi-hypotheses principle, we created a unique proprietary library of CRBN ligands that complements the chemical diversity space covered by commercial libraries (**Fig. 1C**).

### High-throughput global proteomics screening identified a potent and selective TBK1 degrader

High-throughput mass-spectrometric proteomic profiling has emerged as a powerful strategy for neosubstrate-agnostic discovery of MGDs in living cells^42, 51^. Since even weak degraders can provide attractive starting points for optimization, we chose HEK293 cells stably overexpressing CRBN (HEK293-CRBNoe) as a screening system with enhanced detection sensitivity^42, 52^. The screening platform was optimized for both high-throughput and maximum data quality by using automated cell treatment and sample preparation, in a streamlined workflow supporting the screening 40 compounds in duplicate treatments (10 µM, 20 h) with 16 vehicle controls per 96-well plate. The resulting MS samples were analysed using label-free, data-independent acquisition mass spectrometry (DIA-MS^53^), followed by automated statistical data analysis. We screened over 4,000 compounds from the MGL.v2 on this platform, corresponding to ∼10,000 single-shot DIA-MS samples, with an average proteome depth of ∼10,500 quantified proteins per run. From this dataset we identified CCT412020, which selectively reduced TBK1 abundance by more than 4-fold (log_2_FC = −2.03; **Fig. 2A**). The effect was found to be time-dependent, with a log_2_FC(TBK1) value of −1.36 after 6 hours (**Suppl. Fig. 1A**). Interestingly, CCT412020 also modestly decreased levels of the TBK1-interacting proteins TANK and AZI2 (20 h, log_2_FC = −0.41 and −0.32, respectively, **Fig. 2A**), consistent with possible bystander degradation. We also detected several weaker degraders displaying less than two-fold TBK1 downregulation, underscoring the sensitivity and dynamic range of the screening platform. Importantly, CCT412020 also induced strong TBK1 degradation in parental HEK293 cells (**Fig. 2B**). In contrast, TBK1 downregulation was abolished in CRBN-knockout cells (**Fig. 2C**), confirming that CCT412020 acts through a CRBN-dependent mechanism.

**Figure 2.**
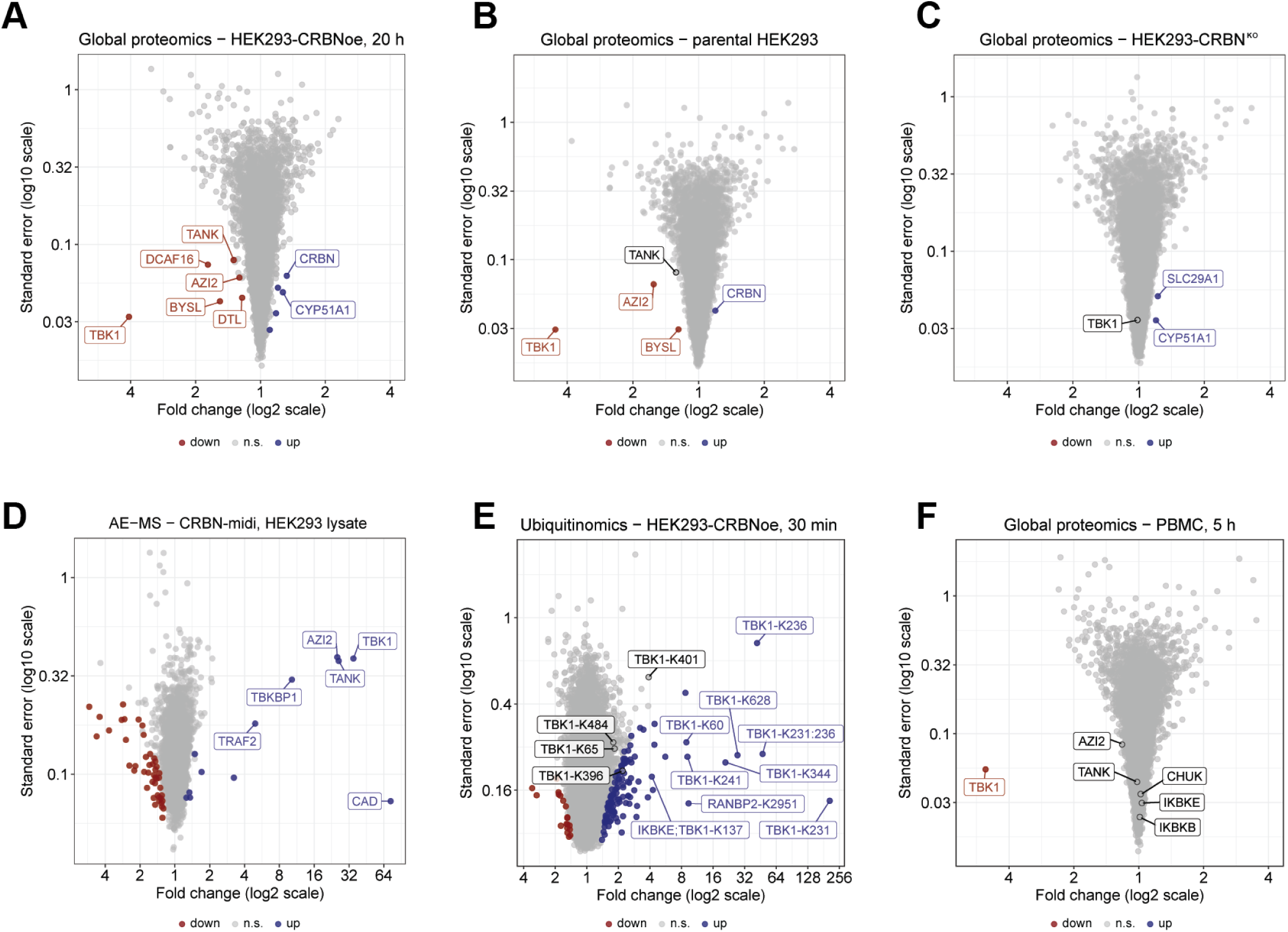
Identification and characterization of a TBK1 molecular glue degrader using high-throughput proteomics. Volcano plots show fold-changes relative to vehicle (x-axis, log_2_ scale) and standard errors (y-axis, log_10_ scale). Up-and downregulated proteins (q < 0.01) or ubiquitination sites (q < 0.05) are coloured in blue and red, respectively. Not significant (n.s.) regulations are coloured grey. **(A)** Global proteomic profile of screening hit CCT412020 (10 µM, 20 h) in HEK293-CRBNoe cells identified from high-throughput proteomic screening of around 4,000 MGD candidates. **(B)** Global proteomics analysis demonstrating TBK1 degradation activity of CCT412020 in parental HEK293 cells (10 µM, 20 h). **(C)** Absence of TBK1 degradation upon CCT412020 treatment (10 µM, 20 h) in HEK293-CRBN^KO^ cells. **(D)** AE-MS of CRBN-interacting proteins from HEK293 cell lysates captured by immobilised CRBN^midi^ bait protein upon addition of 10 µM CCT412020. **(E)** Global ubiquitinomics by K-GG remnant profiling showing induced ubiquitination of multiple TBK1 lysine residues upon 30 min CCT412020 treatment (10 µM) in HEK293-CRBNoe cells. **(F)** Global proteomics analysis demonstrating selective TBK1 degradation in PBMCs upon CCT412020 treatment (10 µM, 5 h).

To investigate if CCT412020-induced degradation results from a direct interaction between TBK1 and CRBN, we performed interactomics profiling using affinity enrichment mass spectrometry (AE-MS). Similar to a recently described approach^43, 54^, we spiked HEK293 lysates with biotinylated CRBNmidi^55^ to capture CRBN-associated proteins in presence of CCT412020 (**Fig. 2D and Suppl. Fig. 1B**). The AE-MS data analysis revealed a robust enrichment of TBK1 (log_2_FC = 5.13), together with weaker enrichments of known TBK1-associated proteins such as AZI2, TANK, TKBKP1, and TRAF2^56^. This pattern is consistent with the direct recruitment of TBK1 to CRBN by CCT412020, along with co-enrichment of the native complex partners of TBK1. The only additional, strongly enriched protein was CAD, which has been reported as a potential TRAF2 interactor^57^. For additional mechanistic validation, we assessed CCT412020-promoted ubiquitination using global KGG remnant profiling in HEK293-*CRBN^oe^* cells^58, 59^. This analysis revealed extensive CCT412020-induced ubiquitination of TBK1, with prominent modification at K231 and K236 (**Fig. 2E and Suppl. Fig. 1C**), consistent with CRBN-mediated Ubiquitin tagging of TBK1 preceding proteasomal degradation.

To demonstrate the translatability of selective CCT412020-induced TBK1 degradation to a native context, we performed unbiased global proteomics of peripheral blood mononuclear cells (PBMCs). Gratifyingly, in this primary cell context the compound also produced robust and selective TBK1 protein depletion (**Fig. 1F and Suppl. Fig. 1D**). This was particularly encouraging as in contrast to HEK293 cells, PBMCs express the TBK1 paralogue IKKε, which high sequence homology presents a considerable challenge for the development of TBK1-selective inhibitors. Notably, CCT412020 showed excellent selectivity over IKKε and related kinases, such as IKKα (encoded by CHUK) and IKKβ. Global ubiquitinomics in PBMCs again highlighted lysines 231 and 236 as the most strongly induced TBK1 ubiquitination sites, supporting the validation of TBK1 as a novel CRBN neosubstrate in a physiologically relevant context primary cell context (**Suppl. Fig. 1E and F**).

### Orthogonal studies confirmed CCT412020-induced potent, selective and CRBN-dependent TBK1 degradation

Next, we further corroborated CCT412020-driven TBK1 degradation by immunoblotting in wild-type HEK293 cells. Incubation of HEK293 cells with CCT412020 over 24 hours resulted in 80% loss of TBK1 protein (**Fig. 3A**). To interrogate the pathway requirements, we assessed CCT412020-induced TBK1 degradation in the presence of the proteasome inhibitor bortezomib (BTZ), the neddylation inhibitor MLN-4924 (as NEDD8 is required for CRL4^CRBN^ Cullin-Ring E3 ligase activity), and in a CRBN-knockout HEK293 line^60^. In each setting, CCT412020 failed to reduce TBK1 protein levels, consistent with a mechanism requiring CRBN, an active Cullin-RING E3 ligase, and the proteasome (**Fig. 3B, C**). Co-treatment with the CRBN ligand lenalidomide markedly suppressed TBK1 degradation, shifting the DC_50_ from 10 nM to 1 µM, which further supports CRBN engagement. In contrast, the TBK1 kinase inhibitor GSK8612^61^ did not compete with CCT412020, consistent with a non-catalytic binding mode and a distinct binding site (**Fig. 3D**). Collectively, these data confirm that CCT412020 induces TBK1 degradation through MGD-like CRBN-dependent Ubiquitin-proteasome pathway activity (**Fig. 3B, C, D**). Importantly, consistent with the proteomics profiling, CCT412020 did not affect the closely related kinases IKKε, IKKα and IKKβ, nor did it alter levels of other pathway components, such as RIPK1 and TAK1 in HeLa cells (**Fig. 3E**).

**Figure 3.**
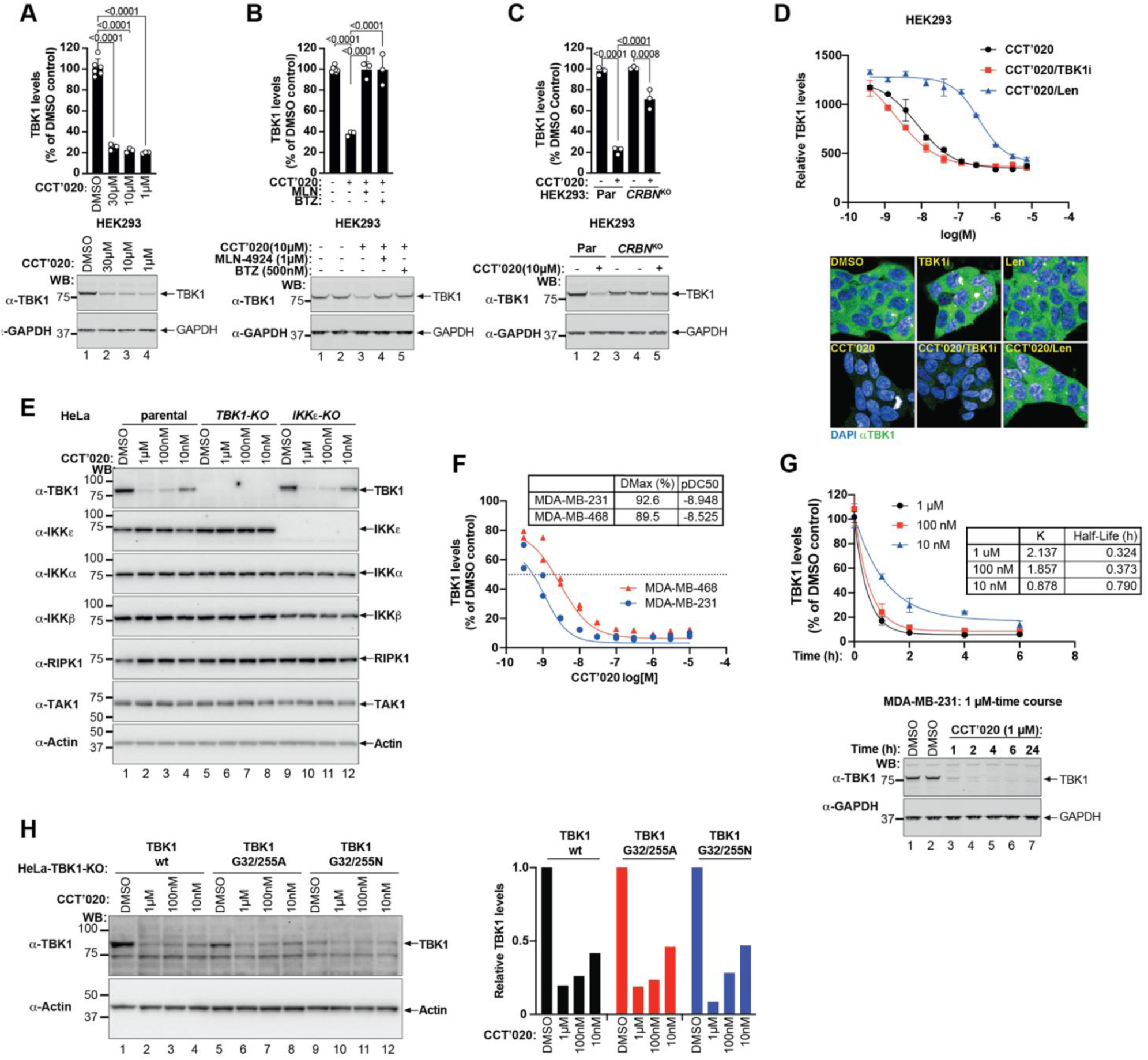
Orthogonal validation of the TBK1 molecular glue degrader, CCT412020. **(A)** Western blot analysis HEK293 cells treated with indicated concentrations of CCT412020 for 24 h. TBK1 abundance relative to GAPDH was quantified by densitometry and normalised to vehicle control (above). **(B)** Western blot analysis of HEK293 cells pretreated for 1h with bortezomib (BTZ, 500 nM) or MLN-4924 (1 µM) before CCT412020 (10 µM) exposure (6 h). TBK1 abundance relative to GAPDH was quantified by densitometry and normalised to vehicle control (above). **(C)** Western blot analysis of HEK293 parental or CRBN-knockout (CRBN^KO^) cells treated with CCT412020 (10µM) for 24 h. TBK1 abundance relative to GAPDH was quantified by densitometry and normalised to vehicle control (above). **(D)** Micro-confocal images of TBK1 endogenous levels in HEK293 cells pretreated with TBK1i (GSK8612, 1 µM, 10 min) or lenalidomide (20 µM, 10 min) and followed by indicated concentrations of CCT412020 (20 h). Above quantification of relative TBK1 levels. **(E)** Western blot analysis of parental HeLa, HeLa-TBK1-KO and HeLa-IKKε-KO cells treated with indicated concentrations of CCT412020 (18 h). TBK1 and relevant kinases levels were visualised. **(F)** CCT412020 potently degrades TBK1 in breast cancer cell lines MDA-MB-231 (pDC_50_ 8.9, D_max_ 93%) and MDA-MB-468 (pDC_50_ 8.5, D_max_ 90%). TBK1 abundance relative to GAPDH was quantified by densitometry and normalized to vehicle control. **(G)** Degradation time-course in MDA-MB-231 cells demonstrating fast degradation kinetics resulting in nearly complete degradation within 1 h. **(H)** Western blot analysis of HeLa-TBK1-KO cells reconstituted with TBK1 wild-type, TBK1G32/255A or TBK1G32/255N. Cells were treated with indicated concentrations of CCT412020 (18 h). TBK1 abundance relative to β-actin was quantified by densitometry and normalised to vehicle control.

We next determined the CCT412020 degradation profile across a panel of breast cancer cell lines, including MDA-MB-231, MDA-MB-468, BT549 and MCF7. In this panel, CCT412020 demonstrated strong, low nanomolar degradation potency and rapid kinetics, in most examples achieving near complete depletion of TBK1 within one to four hours (**Fig. 3F, G, and Suppl. Fig. 2A, B, C, D**). As seen previously (**Fig. 3E**), we did not observe degradation of IKKε, in BT-549 and MCF7 cells, but rather a slight increase in IKKε levels in MCF7 cells (**Suppl. Fig. 2E**).

As previously reported, CRBN-based molecular glues show limited activity in murine cells due to species-specific differences in CRBN^34^. Prior studies have demonstrated that introducing human CRBN, or a ‘humanised’ mouse CRBN variant (I391V), can restore glue-dependent degradation^34^. To enable assessment in a murine tumour model, we generated a MC38 mouse colon cancer line stably expressing human CRBN (MC38*^hCRBN^*). In contrast to parental MC38 cells, CCT412020 treatment of MC38*^hCRBN^*cells induced degradation of endogenous mouse TBK1, confirming productive engagement of the CRBN-dependent degradation machinery in this setting (**Supp. Fig. 2F**).

### Mutations in the predicted G-loop did not rescue CCT412020-induced TBK1 degradation

A computational survey of TBK1 structures in the Protein Data Bank (PDB) and AlphaFold models identified two putative G-loop regions within TBK1. The first motif includes residues 27-33 (RHKKTG^32^D) and closely resembles the CK1α G-loop (Cα RMSD = 0.4 Å, **Suppl. Fig. 2G**)^31, 33, 38, 62^. The second motif encompasses residues 250-256 (QKAENG^255^P) and shows substantially lower structural similarity to CK1α (Cα RMSD = 1.95 Å, **Suppl. Fig. 2H**). Previous studies have demonstrated that that mutation of the G-loop conserved glycine can abolish neosubstrate degradation by CRBN-based MGD^38^. To assess whether CCT412020-induced TBK1 degradation occurs through a similar G-loop-dependent mechanism, we generated TBK1 double mutants in which G32 and G255 in the predicted G-loop regions were replaced with either alanine or asparagine. Wild-type and mutant TBK1 constructs were reconstituted in HeLa TBK1-knockout cells, treated with increasing concentrations of CCT412020, and TBK1 abundance was quantified by Western blotting. Strikingly, both TBK1 double mutants were degraded with efficiencies comparable to wild-type TBK1, indicating that disruption of these predicted G-loop motifs does not impair CCT412020-mediated TBK1 degradation and suggesting potential non-canonical interactions with CRBN (**Fig. 3H**). To determine unambiguously whether TBK1 engages directly with the CRBN/CCT412020 complex, we performed analytical size-exclusion chromatography experiments with purified proteins (**Suppl. Fig. S3**). We found that in presence of CCT412020 TBK1 co-eluted with the CRBN/DDB1 complex at an earlier retention volume than in absence of CCT412020, demonstrating that a direct complex between CRBN/DDB1 and TBK1 was formed in the presence of the MGD.

### Cryo-EM reveals CCT412020 in a non-canonical complex with TBK1 homodimer and CRBN

To understand the mechanism of action of CCT412020 at the molecular level, we determined a cryo-EM structure of TBK1 bound to the Δ39-CRBN/ΔBPB-DDB1 complex in presence of the molecular glue CCT412020. The structure obtained at 3.3 Å resolution revealed CCT412020 bound in complex with a TBK1 dimer and CRBN/DDB1 (**Fig. 4A**). As expected, the glutarimide moiety of CCT412020 was bound in the tri-tryptophan pocket of CRBN. However, unexpectedly, the rest of the molecule was found extending away from the CRBN surface towards the interface of a TBK homodimer, where it intercalates between the predicted G-loop of one monomer and an α-helix in the other monomer. The presence of a TBK1 dimer resulted in two available CRBN/DDB1 binding interfaces. Indeed, two particle populations were observed during processing which contained either one or two copies of the CRBN/DDB1 complex bound to opposite faces of the TBK1 dimer (**Suppl. Fig. 4A**). The structure revealed a novel TBK1 degron binding mode in which the predicted RHKKTG^32^D G-loop is located on the opposite side of the compound, as compared to a canonical CRBN neosubstrate G-loop degron. Additionally, no interaction was observed between CCT412020 and the predicted G-loop glycine 32. This is in stark contrast to other well characterised neosubstrates, such as CK1α, for which the interaction of the G-loop glycine with the IMiD is essential for recruitment of the G-loop degron to CRBN and subsequent degradation of the neosubstrate. These structural differences explain why the TBK1 G32A and G32N mutations did not abolish TBK1 degradation as they do for other canonical neosubstrates. Based on the structure of the TBK1/CCT412020/Δ39-CRBN/ΔBPB-DDB1 complex, we designed a new set of TBK1 mutants containing bulky side chains at three positions around the CCT412020 binding site (L8W, G32W and N578W), aimed at disrupting TBK1-CCT412020 interactions by steric hindrance. Indeed, all three mutants prevented TBK1 degradation by CCT412020 in HeLa cells, confirming that the observed cryo-EM complex occurs in the living cells, too (**Fig. 4B**).

**Figure 4.**
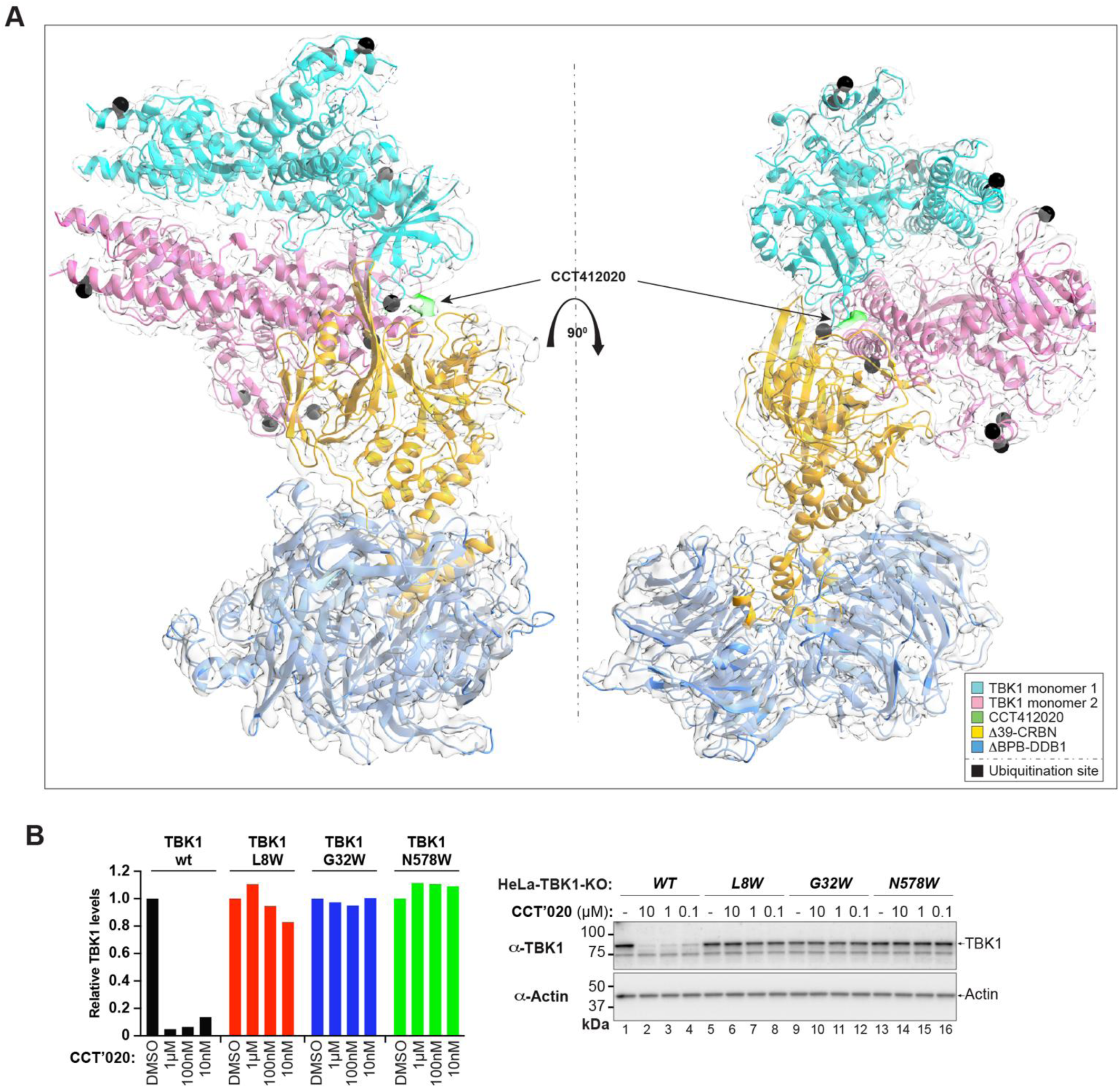
CCT412020 promotes the formation of a ternary complex between TBK1 and CRBN/DDB1, leading to TBK1 ubiquitination. **(A)** The TBK1 dimer (cyan and pink), CRBN (yellow) and DDB1 (light blue) are represented as cartoon traces. The EM map is represented as a semi-transparent grey surface in panel A, with the density corresponding to CCT412020 highlighted in green. Lysine residues identified as sites of ubiquitination are shown by black spheres. Structural figures were generated using ChimeraX. **(B)** Western blot analysis of HeLa-TBK1-KO cells reconstituted with TBK1 wild-type, TBK1-L8W, TBK1-G32W or TBK1-N578W. Cells were treated with indicated concentrations of CCT412020 (18 h). TBK1 abundance relative to β-actin was quantified by densitometry and normalised to vehicle control.

### CCT412020-induced TBK1 depletion sensitises cancer cells to TNF- and IFN-mediated cytotoxicity

TBK1 integrates innate immune signalling downstream of pattern-recognition receptors and shapes cytokine-driven cell fate decisions^8^. To place TBK1 degradation in this signalling context, we first examined its role in TLR-dependent pathway activation. TBK1 is a central mediator of IRF3 activation downstream of toll-like receptor (TLR3) signalling^10, 12, 63^. Accordingly, we treated HEK293-Dual™ hTLR3 reporter cells with poly(I:C) in the presence or absence of CCT412020. Degradation of TBK1 with CCT412020 potently suppressed IRF3 pathway activation (**Fig. 5B**). As expected, NF-κB pathway activation was not inhibited under these conditions (**Fig. 5C**). We next evaluated the role of TBK1 in TNF-signalling, where TBK1 has also been proposed to act as a modulator^7, 10, 12^. Consistent with previous reports indicating that loss of TBK1 can enhance NF-κB activation, CCT412020 increased NF-κB signalling (**Fig. 5D**)^64^.

**Figure 5.**
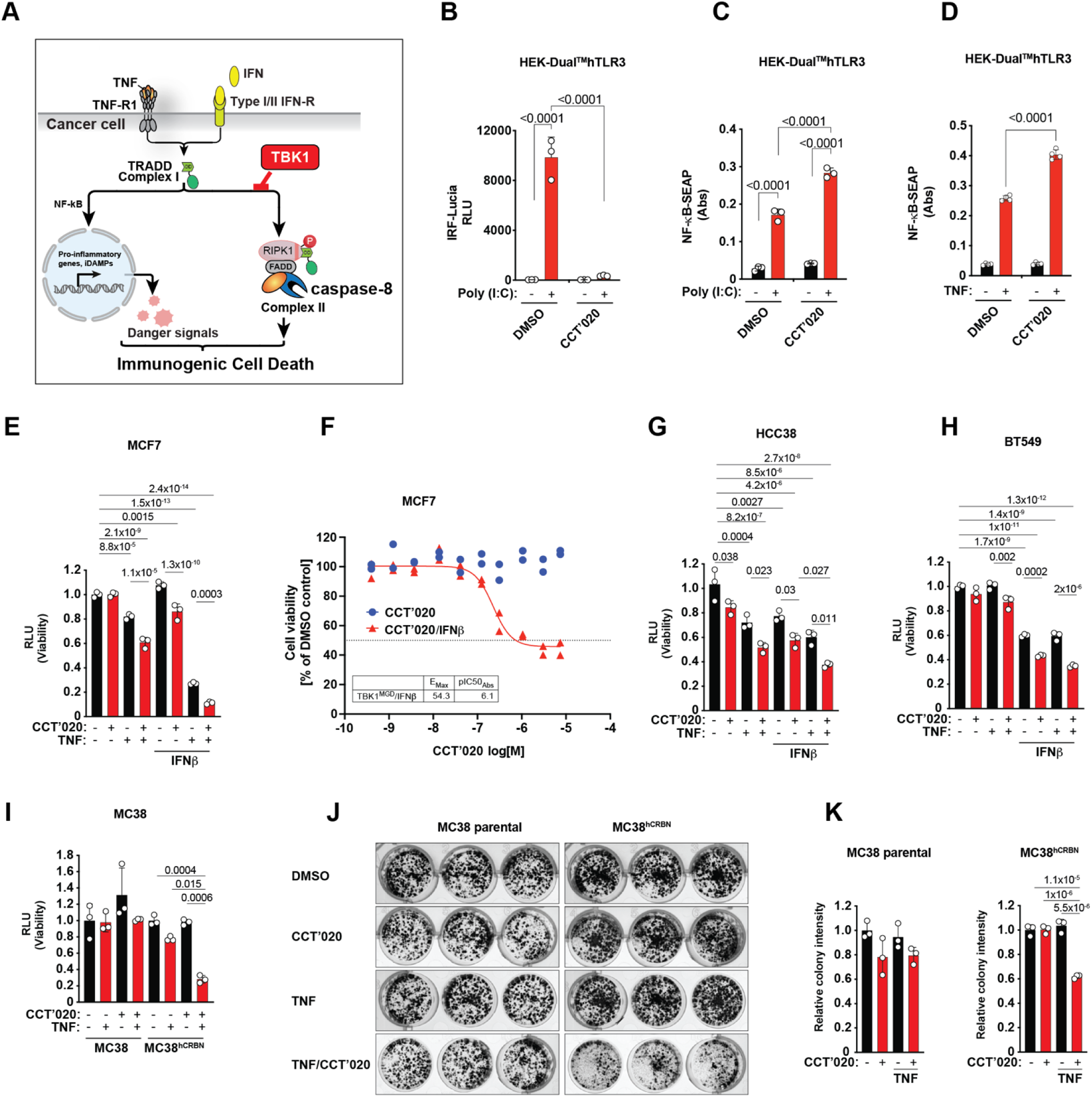
Targeting TBK1 for degradation sensitises breast cancer cells to TNF and IFN induced cell death. **(A)** Schematic representation depicting the role of TBK1 in regulation of TNFR1- and IFN-induced signalling and cell death. **(B)** IRF pathway activation was assessed using a secreted Lucia luciferase reporter assay in HEK-Dual^TM^hTLR3 cells. Cells were preincubated with 1 µM CCT412020 (12 h) prior to stimulation with poly (I:C) (1 µg/ml) for 6 h. IRF-dependent signalling was quantified by measuring secreted Lucia luciferase activity in the culture supernatant using a luminometer. **(C)** NF-κB pathway activity was measured using a SEAP reporter assay in HEK-Dual™ hTLR3 cells. Cells were pre-incubated with the 1 µM CCT412020 for 12 h prior to stimulation with poly(I:C) (1 µg/mL) for 6 h. NF-κB-dependent SEAP activity was quantified by measuring absorbance at 595 nm. **(D)** TNF induced NF-κB pathway activity was measured using a SEAP reporter assay in HEK-Dual™ hTLR3 cells. Cells were pre-incubated with the 1 µM CCT412020 for 12 h prior to stimulation with TNF (10 ng/mL) for 6 h. NF-κB–dependent SEAP activity was quantified by measuring absorbance at 595 nm. **(E)** Cell viability was assessed using a CellTiter-Glo (CTG) assay in the breast cancer cell lines MCF7. Cells were pre-treated with CCT412020 (1 µM) and IFNβ (1 ng/ml) for 24 h, after which TNF (10 ng/ml) was added either as a single agents or in combination for additional 44 h. Viability was quantified following treatment to evaluate the impact of TBK1 degradation alone or in combination with pro-inflammatory cytokine signalling. **(F)** Dose-response analysis of CCT412020 in MCF7 cells in the presence or absence of IFNβ (1 ng/ml) for 48h. Cell viability was assessed using a CellTiter-Glo assay, and DC₅₀ values were determined from a representative experiment. **(G)** Cell viability in HCC38 cells was assessed using a CellTiter-Glo (CTG) as in **(E)**. **(H)** Cell viability in BT549 cells was assessed using a CellTiter-Glo (CTG) as in **(E)**. **(I)** Cell viability was assessed using a CellTiter-Glo (CTG) assay in MC38 and MC38^hCRBN^ cells. Cells were treated with CCT412020 for 18 h in the presence or absence of TNF. Viability was quantified following treatment to evaluate the impact of TBK1 degradation and the contribution of human CRBN expression to TNF-mediated cytotoxic responses. **(J)** Long-term clonogenic survival (7 days) was assessed in MC38 and MC38^hCRBN^ cells following treatment with CCT412020 in the presence or absence of TNFα. Cells were treated as indicated and allowed to grow for colony formation. Representative images of stained colonies in culture wells are shown, and clonogenic survival was quantified **(K)** from scanned plates and plotted as indicated.

Having established that TBK1 degradation differentially tunes innate immune transcriptional outputs, blunting IRF3 while augmenting NF-κB, we next examined the functional consequences for cytokine-induced cytotoxicity downstream of TNFR1 and interferon receptors. Across a panel of human cancer cell lines, CCT412020 sensitised cells to TNF and/or interferon (IFN), resulting in reduced viability in most models tested, consistent with a protective role for TBK1 during cytokine-driven stress (**Fig. 5E, F, G, H and Suppl. Fig. 5A, B**).

We extended these findings to a murine setting using MC38*^hCRBN^*cells that expresses human CRBN. In MC38*^hCRBN^*, but not parental MC38 where the glue is inactive, TBK1 degradation markedly enhanced sensitivity to TNF-induced killing (**Fig. 5 I, J, K**). TBK1 degradation also sensitised cells to necroptotic stimuli in these cells (TNF/emricasan^65^) (**Suppl. Fig. 5C**). Together, these data demonstrate that TBK1 degradation rewires innate immune and cytokine signalling, suppressing IRF3 activation while enhancing NF-κB signalling and sensitizing cancer cells to TNF- and IFN-induced cell death. Importantly, these effects are consistently more pronounced than those achieved by kinase inhibition alone, highlighting distinct functional consequences of TBK1 protein loss.

## Discussion

The clinical success of ICB therapies over the past decade has transformed the treatment of multiple cancer types, establishing immunotherapy as an effective treatment for patients with advanced malignancies who previously had limited options^66^. However, durable benefit is limited to a small subset of patients, largely due to primary or acquired resistance^67^. Emerging evidence indicates that tumour-intrinsic TBK1 activity is an important determinant of resistance to ICB^7, 8, 10, 12^. TBK1 kinase integrates innate immune and death-receptor signalling to tune the balance between immunogenic and non-immunogenic cell death. By blocking caspase-8- and RIPK3-dependent immunogenic cell death pathways, TBK1 acts as a molecular brake on immunogenic cell death and the ensuing anti-tumour immune response^10, 12^. Consistently, genetic ablation of TBK1 sensitises tumour cells to effector cytokines such as TNF and IFNγ, thereby lowering the cytotoxicity threshold for immune-mediated killing, enhances adaptive immunity and improves responses to ICB in preclinical models. Taken together, these findings highlight TBK1 as a compelling therapeutic target to boost anti-tumour immunity and overcome resistance to cancer immunotherapy.

The only approved inhibitor of TBK1 is momelotinib, a multi-kinase inhibitor predominantly targeting JAK1/2, which is used for the treatment of splenomegaly in myelofibrosis patients^68^. Earlier associations of TBK1 activity with autoimmune disorders led to extensive drug discovery efforts across industry and academia that resulted in discovery of potent TBK1 inhibitors, such as BX-795 (IC_50_ = 6 nM), MRT-67307 (28 nM), BAY-985 (2 nM) and A1 (0.775 nM), however, all lack selectivity, especially against the closely related IKKε^69, 70, 71, 72, 73, 74^. More recently, GSK8612 was reported as a highly selective inhibitor of TBK1^61^. While less potent, with a TBK1 IC_50_ value of 158 nM, GSK8612 displayed 63-fold selectivity for the next highest affinity kinase, and two orders of magnitude selectivity over IKKε. In primary immune cells GSK8612 inhibited IRF3 phosphorylation and IFNα secretion with IC_50_ values of around 1 μM^61^. TBK1 degraders have also been described. One of the earliest PROTACs from Arvinas, the VHL-based PROTAC **3i**, is still the most potent TBK1 degrader in the literature, with DC_50_ value of 9 nM^75^. Interestingly, this PROTAC displayed higher potency and selectivity over IKKε than the corresponding inhibitor, although its broader proteome selectivity profile was not disclosed. Another TBK1 PROTAC designed to recruit CRBN, UNC6587, showed only a modest reduction in TBK1 protein levels in clear cell renal cell carcinoma (ccRCC) isogenic cell lines^76^. Recently, a covalent RNF126-directing TBK1 degrader was described as a potential therapeutic approach for autosomal dominant polycystic kidney disease^77^. This compound (**30**) displayed moderate degradation potency with DC_50_= 350.8 nM and D_max_ 91.2% in HEK293T cells, whereas the effect on IKKε or the broader proteome were not disclosed.

Building on recent advances and the promise of MGD drug discovery, we developed a next-generation MGD discovery platform based on high-throughput global proteomic screening of an advanced, rationally designed CRBN-targeting molecular glue library. For the design of our next generation library, MGL.v2, we moved from former strategies solely driven by the desire to maximise structural diversity to a more focused, knowledge-driven multi-hypothesis approach, extensively relying on emerging internal and external data. One example of this focused diversity approach is based on the “degron-targeting” hypothesis, inspired by the observation that MGDs with more extensive interactions with the neosubstrate degron motif tend to display a more potent and selective degradation profile.

High-throughput proteomics screening is another key component our MGD discovery platform. An important advantage of MS-based proteomics screening is unbiased, target- and E3-ligase agnostic nature, enabling simultaneous discovery of novel targets and evaluation of degrader selectivity. In contrast, previous phenotypic screening approaches relied mostly on cell viability as the assay readout, readout, potentially missing weak degraders of relevant cancer targets and even strong degraders of non-essential proteins that may be of therapeutic utility in other diseases. Our high-throughput screening platform combines automated treatment and sample preparation, with DIA-MS data acquisition and fully automated in-house developed data analysis software. This enabled rapid screening of over 4,000 compounds from our MGL.v2, achieving an average depth of more than 10,000 quantified proteins per sample. One of the identified hits was the first-in-class TBK1 molecular glue degrader CCT412020. Extensive proteomics, ubiquitinomics, and pharmacological characterisation confirmed a highly potent, deep and rapid TBK1 degradation profile of CCT412020 in multiple cell lines, including breast cancer cell line MDA-MB-231, with TBK1 DC_50_ value of 1 nM (D_max_ 96%). Interestingly, while TBK1 contains a predicted G-loop degron, the degron G32A and G32N mutations did not rescue TBK1 from the CCT412020-induced degradation. This result suggested that a canonical degron motif is not involved in the CCT412020-driven TBK1 recruitment to CRBN. This observation was confirmed and rationalised by a cryo-EM structure of CCT412020 in complex with CRBN/ΔBPB-DDB1 and a TBK1 homodimer. This structure revealed an unprecedented binding mode with the TBK1 G-loop binding on the opposite side of the MGD CCT412020, which intercalates in an unconventional binding site at the interface between the two TBK1 monomers. The TBK1 depletion induced by CCT412020 triggers immunogenic cell death and pro-inflammatory cytokine release in both human and murine (hCRBN-expressing) cancer cells, providing a tractable strategy to sensitise tumours to immunotherapy^7, 8^.

## METHODS

### Reagents

2-Chloroacetamide (CAA), Tris(2-carboxyethyl)phosphine hydrochloride (TCEP), NaCl, Na_2_HPO_4_, 3-(N-morpholino)propanesulfonic acid (MOPS), Sodium deoxycholate (SDC), Tris(hydroxymethyl)-aminomethane (Tris), trifluoracetic acid (TFA), formic acid (FA), and acetonitrile were from Merck. nUndecyl-β--Maltoside (UDM) from Anatrace. Protease inhibitor mix from ThermoFisher Scientific (A32955). Trypsin from Promega. PTMScan^®^ HS Ubiquitin/SUMO Remnant Motif (K-ε-GG) Kit (#59322) from Cell Signaling Technology. IFNb was Peprotech, 300-02BC and TNF was from Enzo, ALX-522-008-C050, Poly (I:C) (InvivoGen tlrl-pic), MLN4924 (NEDD8 inhibitor, Tocris, 6499/10), bortezomib (Tocris, 7282/5), RIPA buffer (ThermoFisher Scientific, # 89900), Halt Protease and Phosphatase Inhibitor Cocktail (Thermo Fisher Scientific, #78440), Pierce BCA Protein Assay Kit (Thermo Fischer Scientific, #23227), NuPAGE Sample Loading Buffer (ThermoFisher Scientific, #NP0007), 4-12% Bis Tris NuPage gel (Thermo Fisher Scientific, #NP0321), NuPage MOPS running buffer (Thermo Fisher Scientific, #NP0001). Gel electrophoresis system (Bio-Rad, 1645050), iBlot3 system with 0.2 µm nitrocellulose membrane (ThermoFisher IB33001X3) iBind Flex Western Device (ThermoFisher Scientific, #SLF2000). Antibodies used for hit confirmation and mechanism of action Western blots: anti-TBK (Proteintech, #28397-1-AP, 1:1000), anti-GAPDH (Proteintech, #60004-1-IG, 1:20,000), goat anti-mouse and goat anti-rabbit secondary antibodies IRDye® 680RD Goat Anti-Mouse IgG (Licor Biosciences 926-68070), IRDye® 800CW Goat Anti-Rabbit IgG 926-32211. Other WB antibodies: anti-TBK1 (CST #51872), anti-IKKe (CST #3416), anti-IKKe (CST #2905), anti-IKKa (CST #2682), anti-IKKb (CST #2684), anti-RIPK1 (CST #3493), anti-TAK1 (CST #4505), anti-CRBN (CST #71810), anti-Actin (Sigma #A5441). For immunofluorescence assay anti-TBK1 (CST #38066) was used.

### Cell lines

MCF-7, HCC38, BT-549, MDA-MB-231, MDA-MB-468, HeLa, and PANC-1 cell lines were obtained from ATCC. MC38 cells were purchased from Kerafast, CRBN KO cells used for mechanism of action western blots were published previously^78^. All cell lines were cultured in Dulbecco’s Modified Eagle Medium (DMEM) supplemented with 10% foetal bovine serum (FBS; Sigma-Aldrich, #A2153) and 1% penicillin–streptomycin. MC38 cells were additionally cultured in the presence of 5 mM HEPES. All cell lines were maintained at 37°C in a humidified incubator with 10% CO_2_.

### Cell culture for proteomics

Parental HEK293, HEK293 stably overexpressing CRBN (HEK293-CRBNoe)^42^, and HEK293 with CRBN-CRISPR knockout (HEK293-CRBN^KO^) cells were cultured in DMEM (VWR) supplemented with 10% FCS (Thermo Fisher Scientific) (additionally supplemented with puromycin (2 µg/ml) for HEK293-CRBNoe and HEK293-CRBN^KO^ cell culture). Peripheral blood mononuclear cells (PBMCs) were isolated from fresh whole blood (Donas GmbH) using Ficoll-Paque PLUS density gradient media (Cytiva) according to the manufacturer’s protocol. Briefly, one volume of whole blood was diluted with one volume Dulbecco’s PBS (DPBS), then gently added to one volume density gradient media. Following centrifugation (750 ×g, 15 min, r.t.) with gentle braking, the mononuclear cell layer was transferred to a fresh tube, diluted with DPBS, centrifuged (560 ×g, 5 min, r.t.), and aspirated. The cell pellet was washed with DPBS twice, then treated with 1× Erythrocyte Lysis Buffer (EasyLyse, S2364, Dako) for 5 min. Following dilution with DPBS, cells were centrifuged (560 ×g, 5 min, r.t.), aspirate, washed once with DPBS, and once with RPMI +10% FBS. The thus isolated PBMCs were cultured in RPMI +10% FBS and used in proteomic experiments.

### Global proteomics

For high-throughput proteomic screening and global proteomics experiments, cells were cultured in 96-well plates with respective media and treated with the indicated compounds (1000× DMSO stocks) for the indicated time. Cells were lysed with UDM buffer (0.05% w/v UDM, 75 mM Tris-HCl pH 8.5, 40 mM CAA, 10 mM TCEP) and the samples were incubated at 80 °C for 10 min with gentle shaking (400 rpm). The samples were cooled to room temperature and proteins were digested overnight at 37 °C using 400 ng trypsin (Promega) per well. The resulting peptides were desalted using in-house prepared, 200 µL two plug C18 StageTips (3M Empore) ^79^and then analysed by LC-MS/MS.

### Global ubiquitinomics

Global ubiquitinomics experiments were carried out similar to a reported procedure^58^. In brief, cells were cultured in 6- or 12-well plates and treated with the indicated compound (1000× DMSO stocks) for 30 min, followed by lysis with SDC buffer. The protein concentrations were determined using a BCA assay kit (Merck-Millipore) and the proteins were digested overnight at 37 °C using a ratio of protein:trypsin of 100:1. Immunoprecipitation buffer (50 mM MOPS, pH 7.2, 10 mM Na_2_HPO_4_, 50 mM NaCl) was added along with a K-GG antibody-bead conjugate. After incubation for 2 h on a rotor wheel, beads were washed and peptides eluted according to manufacturer’s instructions. The peptide eluate was desalted using in-house prepared, 200 µL two plug C18 StageTips (3M Empore) and then analysed by LC-MS/MS.

### Affinity enrichment-mass spectrometry (AE-MS)

HEK293 cells were lysed on ice in ice-cold NP-40 buffer (0.05% NP-40, 50 mM Tris-HCl pH 7.5, 150 mM NaCl, 5% glycerol), freshly supplemented with protease inhibitors. The lysate was cleared by centrifugation at 20,000 ×g for 10 min (4 °C) and the supernatant transferred to a fresh tube. The protein concentration was determined using a BCA assay kit (Merck-Millipore) and the lysate concentration was adjusted to 1 mg/mL. Biotinylated CRBN^midi^ ^55^(CRELUX, WuXi AppTec) was added to the lysate along with either DMSO or 10 µM CCT412020 and incubated for 1 hour at 4 °C. Biotin affinity capture was used to isolate CRBN^midi^-bound proteins followed by 4 washes with lysis buffer. Proteins were eluted with UDM lysis buffer and digested by adding 100 ng of trypsin per sample (overnight, 37°C). The resulting peptides were desalted using in-house prepared, 200 µL two plug C_18_ StageTips (3M Empore)^79^ and then analysed by LC-MS/MS.

### LC-MS/MS analysis

Peptides were either analysed on mass spectrometers from Bruker (timsTOF HT or timsTOF Ultra 2) or ThermoFisher (Orbitrap Astral). The LC Setup differed for the various sample types and/or mass spectrometers. For global proteomics on timsTOF instruments and global ubiquitinomics the following LC setup was used: Peptides were loaded on 30 cm reverse-phase columns (75 µm inner diameter, packed inhouse with ReproSil Saphir 100 C18 1.5 µm resin [ra115.9e., Dr. Maisch GmbH]) using either a Vanquish™ Neo system (ThermoFisher) or a nanoElute® 2 system (Bruker). The column temperature was maintained at 60 °C using a column oven. The LC flow rate was 300 nL/min and the complete gradient was 50 minutes (global proteomics) or 45 minutes (global ubiquitinomics). The LC setup of the samples measured with a higher throughput (global proteomics samples that were measured on an Orbitrap Astral instrument using a ∼65 samples per day (SPD) method and AE-MS samples that were measured on the timsTOF Ultra 2 using a ∼100 SPD method) differed as follows: a 17 cm column with a 150 µm inner diameter was used, the flow rate was at 2,000 nL/min and the gradient length was 17 minutes (global proteomics) or 9 minutes (AE-MS), respectively. Eluting peptides measured on the timsTOF instruments were analysed using diaPASEF^53^ (global proteomics and AE-MS) or slicePASEF^80^ (global ubiquitinomics) methods as previously described^42^. For the global proteomics data measured on the Orbitrap Astral^81^, a Nanospray Flex™ source was used, the ionization occurred at +2.2 kV, and the ion transfer tube temperature was 280 °C. The Orbitrap scan resolution was 240,000, the RF lens value was 40%, the AGC target was 500%, and the maximum injection time was 3 ms. A data-independent acquisition scheme with 222 isolation windows with widths between 3 and 10 Th covering an m/z range of 320-1231 was used with the following Astral settings: MS2 scan range: 150 - 2,000 m/z, normalized HCD collision energy: 25%, AGC target: 800%, RF lens: 40%, maximum injection time: 3 ms.

### MS raw data processing

MS raw files were analysed using DIA-NN^82^. Global proteomics and AE-MS raw files were processed with v2.1.0, and global ubiquitinomics raw files with v2.2.0. Reviewed UniProt entries (human, SwissProt 10-2022 [9606]) were used as a protein sequence database for DIA-NN searches. One missed cleavage, a maximum of one variable modification (oxidation of methionine), and N-terminal excision of methionine were allowed. Carbamidomethylation of cysteines was set as a fixed modification, and K-GG (UniMod: 121) was added in case of global ubiquitinomics. All data processing were carried out using library-free analysis mode in DIA-NN. “--tims-scan” was added as an additional command in case of global ubiquitinomics.

### Statistical analysis of proteomics data

Statistical analysis was performed as described previously^42^. DIA-NN outputs were further processed with R. Peptide precursor quantifications with missing values in more than 50% of samples, or <33% of the DMSO-treated samples (for global proteomics; <25% of compound-treated samples in case of global ubiquitinomics) were discarded. Protein abundances (for global proteomics and AE-MS) or K-GG peptide abundances (for global ubiquitinomics) were calculated based on precursor ion intensity levels using the MaxLFQ^83^ algorithm as implemented in the DIA-NN R package (https://github.com/vdemichev/diann-rpackage). Completely missing cases in any of the tested conditions were rescued by accepting low-quality precursors (i.e., q-value > 0.01), where possible. K-GG peptide to site mapping was done using reviewed entries of the human UniProt database (SwissProt-, release 102022). The protein (or peptide) intensities were normalized by median scaling and corrected for variance drift over time (if present) using the principal components (derived from principal component analysis) belonging to DMSO samples. Subsequently, protein (or peptide) intensities were subjected to statistical testing with variance and log fold-change moderation built on^84^￼. P-values corrected for multiple testing using the Benjamini-Hochberg method^85^ were used to assess significance in global proteomics (q-value < 0.01), AE-MS (q-value < 0.01), and global ubiquitinomics (q-value < 0.05) experiments. For comparing global proteome and ubiquitinome data, identifications were mapped at the gene level.

### Immunoblot assay

For selectivity analysis of CCT412020 in HEK293 and HEK293 knockout cells, cells were seeded in 12-well plates at a density of 3.5 × 10⁵ cells per well and allowed to adhere overnight. Cells were then treated with CCT412020 at concentrations of 1, 10, or 30 µM for 24 h prior to lysis. To test the ubiquitination and proteasome dependency cells were pre-treated for 1h with 1 µM MLN-4924 (Nedd8 inhibitor), or 500 nM bortezomib followed by 6h treatment with 10 µM compound. To demonstrate CRBN dependency HEK293 KO cells were treated with 10 µM compound for 24h.

Cells were lysed with RIPA buffer supplemented with Halt Protease and Phosphatase Inhibitor Cocktail. Protein levels were quantified using BCA assay Pierce BCA Protein Assay Kit.

20 µg of protein per well was denatured in NuPAGE Sample Loading Buffer containing SDS and dithiothreitol, and loaded onto a 4-12% Bis Tris NuPage gel in NuPage MOPS running buffer (Thermo Fisher Scientific, NP0001). Following SDS-PAGE electrophoresis (150V, 1h, Bio-Rad System) proteins were transferred onto 0.2 µm nitrocellulose membrane using Thermofisher iBlot3 system under 15V for 6 min. Proteins were detected using the iBind Flex Western Device (Thermo Fisher Scientific, SLF2000), using rabbit polyclonal anti-TBK (Proteintech), 28397-1-AP, 1:1000), mouse monoclonal anti-GAPDH antibody (Proteintech, 60004-1-IG, 1:20,000), goat anti-mouse and goat anti-rabbit secondary antibodies, and imaged using Licor Odyssey system.

All HeLa- and MC38-derived stable cell lines were cultured in 12-well plates and lysed directly in Laemmli sample buffer. Lysates were boiled for 3 min, and proteins were subsequently resolved by SDS–PAGE.

### Pharmacological profiling

Cells were plated in 12-well plates (BT-549 at a density of 4 × 10⁵ cells/well and MCF7 - 3 × 10⁵ cells/well) and incubated overnight before compound treatment. For DC_50_ determination 1mM DMSO stock compound solution was diluted 1000x in complete culture medium to obtain the following working concentrations: 10 µM, 3 µM, 1 µM, 300 nM, 100 nM, 30 nM, 10 nM, 3 nM, 1 nM, 300 pM, maintaining 0.1% DMSO concentration and cells were treated for 24 h before WB analysis For degradation kinetics experiments, cells were treated with 1 µM, 100 nM and 10 nM TBK1^MGD^ with 0.1% DMSO for 0.5, 1, 2, 4, or 6 h prior to WB analysis.

### Pharmacological profiling data analysis

Pharmacological profiling data have been analysed using GraphPad Prism v. 10.5.0. Dose-response data have been represented as % of DMSO sample and % of remaining protein has been plotted on a log_10_ scale as a function of compound concentration. DC_50_ and D_max_ values were calculated from a non-linear regression four-parameter curve fit according to the following equation: Y=Bottom + (Top-Bottom)/(1+(IC_50_/X)^HillSlope), as follows: pDC_50_ = -log(IC_50_), pDC_50Abs_ = -(log(X[50])) and D_max_ = 100-Bottom, where X, compound concentration), Y, response, Top and Bottom, plateaus on Y axis, IC_50_ = DC_50_. Note: in figure 3F pDC_50_Abs have been reported rather than pDC50 (curve inflection point).

Degradation kinetics has been determined Exponential One phase decay equation: Y=(Y0 - Plateau)*exp(-K*X) + Plateau, and Half-life=ln(2)/k, where X, Time, Y, % of remaining protein vs DMSO, K, rate constant equal to the reciprocal of the X axis units.

Cell fitness analysis has been performed using four-parameter curve fit to CellTitreGlo data, calculated as % of DMSO signal and plotted on a log10 concentration scale. Cell viability IC_50_ and Emax were calculated in the same manner as DC_50_ and D_max_.

### Immunofluorescence

Cytoplasmic TBK1 protein degradation was quantified using an immunofluorescence-based high-content imaging assay and analysed with Harmony software (PerkinElmer). Briefly, 40 µL of cells cultured in DMEM supplemented with 10% foetal bovine serum (FBS; Thermo Fisher Scientific) were pre-incubated with competitor compounds for 10 min and subsequently seeded into 384-well PhenoPlates (PerkinElmer) preloaded with CCT412020 or DMSO control. Cells were incubated for 20 h at 37 °C in a humidified CO_2_ incubator, fixed with 2% formaldehyde for 15 min at room temperature, and washed with phosphate-buffered saline (PBS) using a Multidrop Combi dispenser (Thermo Fisher Scientific).

Fixed cells were permeabilized for 15 min at room temperature in PBS containing 0.2% Triton X-100 (Thermo Fisher Scientific, #28314), followed by blocking for 1 h in PBS containing 0.5% bovine serum albumin (BSA; Sigma-Aldrich, #A2153). After washing with PBS, cells were incubated overnight at 4 °C in PBS/BSA with rabbit anti-TBK1 primary antibody (Cell Signaling Technology, #38066), together with DAPI (Thermo Fisher Scientific, #D3571) and Phalloidin–Alexa Fluor 633 (Thermo Fisher Scientific, #A22284). Cells were then washed with PBS and incubated for 1 h at room temperature with Alexa Fluor 488–conjugated donkey anti-rabbit secondary antibody (Thermo Fisher Scientific, #A-21206) in PBS/BSA. Following final washes with PBS, plates were imaged using an Opera Phenix Plus high-content imaging system (PerkinElmer).

### Identification of G-loop Motifs

G-loop degron motifs were identified in structures from the Protein Data Bank (PDB, accessed on 09/25) and the AlphaFold Structure Database (AFDB; version 6) using an in-house computational pipeline. Protein structures were first decomposed into their constituent monomers, and the Define Secondary Structure in Proteins algorithm (DSSP; version 2.3.0) was applied to assign secondary structure elements, solvent accessibility, and intramolecular hydrogen bonds.

A sliding-window approach was then used to analyse G-loop motifs, defined as glycine-containing 7-mers spanning positions G_−5_ to G_+1_. Candidate 7-mers were filtered using the following criteria, derived from known neosubstrate G-loops observed in ternary complex structures: (i) a backbone hydrogen bond between positions G_−4_ and G_+1_ (ΔG_H-bond_ < -1 kJ mol^-1^), (ii) no more than 2 of 7 residues adopting helical secondary structure, and (iii) a minimum solvent accessibility of 30 Å² for G_0_, with solvent accessibility ≥ 35 Å² for at least 4 of 6 residues in the motif.

Compatible motifs were then structurally aligned to the G-loop degron from CK1α (PDB 5FQD) by superimposing Cα atoms of the motif residues, and alignments with a root mean square deviation (RMSD) >2 Å were discarded. Following alignment, steric clashes between backbone atoms in CRBN (PDB 5FQD, chain B) and the surface of the target protein (having residue solvent accessibility > 5 Å) were calculated. Targets exhibiting steric clashes within the degron motif or within 5 Å of this region were removed from further analysis.

### CRBN/DDB1 and TBK1 expression

The human His_6_-ZZ-HRV-3C-CRBN_39-442_ and ΔBPB-DDB1-Strep protein complex and the human His_6_-GST-HRV-3C-TBK1_1-657_ protein were separately expressed in sf9 insect cells using standard protocols. Baculoviruses were generated using the Bac-to-Bac-Baculovirus Expression System (Thermo Fisher Scientific) according to the manufacturer’s instructions. Sf9 cells growing at 27 °C in shaker flasks with Sf-900™ III SFM media were infected with 20 μL virus/10^7^ cells and harvested 72 hours post infection. Cells were harvested by centrifugation at 6200 x g for 20 minutes at 4 °C, and cell pellets were stored at -70 °C.

### Agnostic Ikaros peptide expression

Human Ikaros_140-196_ Q146A G151N with an N-terminal His_6_-MBP-TEV-tag^47^ in a pET28a vector was purchased from Twist Biosciences. Expression was performed using BL21-AI *E. coli* cells grown in Terrific Broth media supplemented with 50 μg/mL Kanamycin and 150 µM Zinc acetate. Cultures were grown at 37 °C until an OD_600 nm_ > 1.0 and protein expression was induced with 0.2 % L-Arabinose and 0.2 mM IPTG. Expression was carried out for 18 hours at 18 °C. Cells were harvested by centrifugation at 6200 x g for 20 minutes at 4 °C, and cell pellets were stored at -70 °C.

### CRBN/DDB1 purification

Cells were re-suspended in buffer 1A (50 mM HEPES pH 8.0, 500 mM NaCl, 0.5 mM TCEP) supplemented with 1 mM MgCl_2_, 1x cOmplete^TM^ ULTRA protease inhibitors and 12.5 U/mL Benzonaze. Cells were lysed by sonication followed by centrifugation at 55,900 x g for 1 hour at 4°C and filtered through a 1.2 μm syringe filter. Clarified lysate was loaded onto a 5 mL HisTrap FF column, washed with buffer 1A supplemented with 10 mM imidazole and eluted with buffer 1B (50 mM HEPES pH 8.0, 500 mM NaCl, 0.5 mM TCEP, 250 mM imidazole). The NaCl concentration was reduced to 100 mM by dilution with buffer 1C (50 mM HEPES pH 8.0, 0.5 mM TCEP) and applied to a 6 mL Resource Q column and eluted with a gradient from 50 to 500 mM NaCl over 10 CVs with buffer 1D (50 mM HEPES pH 8.0, 1 M NaCl, 0.5 mM TCEP). The His_6_-ZZ-tag on CRBN was cleaved by the addition 20 U/mg of HRV-3C protease for >8 hours at 4 °C. NaCl concentration was reduced to 100 mM and cleaved CRBN_39-442_/ΔBPB-DDB1-Strep complex applied to a 6 mL Resource Q column. The bound protein of interest was eluted with a gradient from 50 to 500 mM NaCl over 10 CVs. The protein was further purified using a HiLoad Superdex 200 26/600 pg column pre-equilibrated in a buffer 1E (50 mM HEPES pH 8.0, 200 mM NaCl, 1 mM TCEP). The final protein complex was concentrated to 20 mg/mL using a centrifugal concentrator with a 30 kDa molecular weight cut-off and stored at -70 °C. Final purity was assessed by SDS-PAGE and molar mass by high-resolution intact mass spectrometry.

### TBK1 purification

Cells were re-suspended in buffer 2A (20 mM HEPES pH 7.5, 500 mM NaCl, 0.5 mM TCEP) supplemented with 1 mM MgCl_2_, 1x cOmplete^TM^ ULTRA protease inhibitors and 12.5 U/mL Benzonaze. Cells were lysed by sonication followed by centrifugation at 55,900 x g for 1 hour at 4°C and filtered through a 1.2 μm syringe filter. Clarified lysate was loaded onto a 5 mL HiTrap TALON Crude column, washed with buffer 2A and eluted with buffer 2B (20 mM HEPES pH 7.5, 500 mM NaCl, 0.5 mM TCEP, 150 mM imidazole). The protein was subsequently applied to a 5 mL GSTrap HP column, pre-equilibrated in buffer 2A, and eluted with buffer 2A supplemented with 20 mM Glutathione. The His_6_-GST tag was cleaved by the addition 20 U/mg of HRV-3C protease for >8 hours at 4 °C. Cleaved TBK1_1-657_ was concentrated to 2.5 mL using a centrifugal concentrator with a 30 kDa molecular weight cut-off and further purified using a HiLoad Superdex 200 16/600 pg column pre-equilibrated in a buffer 2C (20 mM HEPES pH 7.5, 300mM NaCl, 1 mM TCEP). Finally, the protein was applied to a 5 mL GSTrap HP column, pre-equilibrated in buffer 2C, and the flowthrough collected. TBK1_1-657_ was concentrated to 2.8 mg/mL using a centrifugal concentrator with a 30 kDa molecular weight cut-off and stored at -70 °C. Final purity was assessed by SDS-PAGE and molar mass by high-resolution intact mass spectrometry.

### Agnostic Ikaros peptide purification

Cell pellets were re-suspended in buffer 3A (50 mM HEPES pH 7.5, 200 mM NaCl, 0.5 mM TCEP, 10% glycerol, 150 µM Zinc Acetate) supplemented with 1 mM MgCl_2_, 1x cOmplete^TM^ ULTRA protease inhibitors and 12.5 U/mL Benzonase. Cells were lysed by sonication followed by centrifugation at 55,900 x g for 1 hour at 4 °C and filtered through a 1.2 μm syringe filter. Clarified lysate was loaded onto a 5 mL MBPTrap, washed with 20 CV of buffer 3A and eluted with buffer 3B (50 mM HEPES pH 7.5, 200 mM NaCl, 0.5 mM TCEP, 10% glycerol, 10 mM Maltose, 150 µM Zinc Acetate). The His_6_-MBP tag was cleaved by incubating with 1:50 TEV protease for >8 hours at 4 °C. Cleaved Ikaros_140-196_ Q146A G151N was applied to a 5 mL HisTrap FF column pre-equilibrated in buffer 3A, and eluted in buffer 3C (50 mM HEPES pH 7.5, 200 mM NaCl, 0.5 mM TCEP, 150 µM Zinc Acetate, 10% glycerol, 250 mM Imidazole). The protein was concentrated to 3.5 mL using a centrifugal concentrator with a 3 kDa molecular weight cut-off and further purified using a HiLoad Superdex 75 26/600 pg column pre-equilibrated in buffer 3D (10 mM HEPES pH 7.0, 240 mM NaCl, 1 mM TCEP, 50 µM Zinc Acetate). The final sample was concentrated to 0.4 mg/mL using a centrifugal concentrator with a 3 kDa molecular weight cut-off and stored at −70 °C. Final purity was assessed by SDS-PAGE and molar mass by high-resolution intact mass spectrometry.

### Cryo-EM grid preparation/ Data collection/ Data processing

The Δ39-CRBN/ΔBPB-DDB1/CCT412020/TBK1 complex was formed by mixing Δ39-CRBN/ΔBPB-DDB1 with CCT412020 and TBK1 in a 1:10:1.2 ratio (10.5 µM Δ39-CRBN/ΔBPB-DDB1:105 µM CCT412020:12.6 µM TBK1). The 20 mg/mL Δ39-CRBN/ΔBPB-DDB1 and 2.8 mg/mL TBK1 stocks were diluted in buffer 4A (20 mM HEPES pH 7.0, 150 mM NaCl, 3 mM TCEP), giving a final DMSO concentration of 1.05%. The complex was incubated on ice for 1 hour and centrifuged at 21,000 x g for 10 minutes at 4 °C before grid preparation. A 10 µM solution of Ikaros_140-196_ Q146A G151N was prepared by diluting the 0.4 mg/mL stock in buffer A and centrifuged at 21,000 x g for 10 minutes at 4 °C before grid preparation.

UltrAuFoil R1.2/1.3 gold grids were plasma cleaned for 50 seconds using a Tergeo plasma cleaner in an air mix at 15 W. A Vitrobot Mark IV was used for cryoEM grid preparation set at 100 % humidity and 4 °C. Grids were pre-treated by the addition of 4 µL of the 10 µM Ikaros_140-196_ Q146A G151N, incubated for 60 seconds, then blotted for 4 seconds, blot force 0. During this incubation time the Δ39-CRBN/ΔBPB-DDB1/CCT412020/TBK1 complex was diluted 10-fold with buffer A to give final concentrations of 1.05 µM, 10.5 µM and 1.26 µM for Δ39-CRBN/ΔBPB-DDB1, CCT412020 and TBK1 respectively. During this incubation time the Δ39-CRBN/ΔBPB-DDB1/CCT412020/TBK1 complex was diluted 10-fold with buffer A to give final concentrations of 1.05 µM, 10.5 µM and 1.26 µM for Δ39-CRBN/ΔBPB-DDB1, CCT412020 and TBK1, respectively. This sample was immediately applied to the grid after the Ikaros_140-196_ Q146A G151N blotting, blotted for 4 seconds, blot force 0, and flash frozen in liquid ethane. Grids were subsequently clipped into autogrid cartridges for use in Thermo Fisher microscope autoloader systems.

Datasets were collected with EPU on a Thermo Scientific Glacios microscope using an acceleration voltage of 200 kV and at a nominal magnification of 165,000 times to give a nominal pixel size of 0.7 Å. Movies were recorded on a Falcon 4i direct electron detector, equipped with a Selectris energy filter set at a slit width of 10 e^-^V. A total electron dose of 60 e-/Å^2^, from a 2.4 second exposure, was collected with a nominal defocus range of -0.8 μm to -1.8 µm. Initially 10,000 movies were collected from one grid. An additional 14,000 movies were subsequently collected with the same settings from a second grid at a tilt angle of 30 degrees to reduce preferred orientation.

Processing was performed in CryoSPARC v4.7.0^86^. Patch motion correction followed by Patch CTF estimation was performed using standard parameters and micrographs were subsequently filtered for excessive motion and poor CTF fits. Particle picking initially using Blob Picker, extracting particles in a box size of 448 x 448 pixels, downscaled to 112 x 112 pixels (pixel size of 2.8 Å) was used to generate initial 2D classes. Two rounds of 2D classification were performed and a subset of the best particles from selected 2D classes were then used to train Topaz^87^ which was subsequently used for particle picking, followed by two rounds of 2D classification. The best particles were subsequently re-extracted in a box size of 512 x 512 pixels, downscaled to 384 x 384 pixels (pixel size of 0.93 Å) and were used to generate 3 ab-initio classes followed by a round of Heterogenous refinement Particles from the best class were taken through to a round of non-uniform refinement, with ‘Optimise per-particle defocus’ enabled, followed by 3D classification into four classes, using a ‘Filter resolution’ of 4 Å. A final round of Non-uniform refinement generated a map at 3.3 Å resolution consisting of 280,220 particles. 3DFSC^88^ analysis showed that the generated map is isotropic with a sphericity value of 0.961. A second map was generated at 3.3 Å resolution consisting of 261,476 particles that was consistent with having two CRBN/DDB1 complexes bound to the TBK1 dimer. Atomic model building and refinement was performed using a previously in-house determined structure of CRBN/DDB1 and the published structure of TBK1 PDB: 6NT9^89^ as starting models. Model building was performed iteratively in Coot 0.9.8^90^ and PHENIX real-space refinement^91^ and the quality of the structure assess with MOLPROBITY^92^. CCT412020 ligand restraints were generated in GRADE^93^. Structures were visualised and figures generated in UCSF ChimeraX^94^. Final model statistics are available in **Suppl. Fig. 4B**.

### Size Exclusion Chromatography assessment of ternary complex formation

Two 60 µL samples of 20 µM Δ39-CRBN/ΔBPB-DDB1 with 25 µM TBK1 were prepared in buffer A (20 mM HEPES pH 7.0, 200 mM NaCl, 1 mM TCEP) and incubated with either 100 µM CCT412020 or the equivalent concentration of DMSO (1%) for 1 hour on ice. Samples were centrifuged at 21,100 x g for 10 minutes before injecting onto a Superdex 200 5/150 GL column, preequilibrated in buffer A, via a 50 µL capillary loop. The column was run at 0.15 mL/minute for 1.5 CV and 50 µL fractions collected. Fractions were analysed by NuPAGE Bis-Tris 4-12% Mini Protein Gels.

### Constructs and stable cell lines

Human TBK1 (wild-type and mutant) and human CRBN GeneArt synthetic constructs (Thermo Fisher Scientific) were subcloned into an in-house PiggyBac expression vector (pGMBE). The resulting constructs were co-transfected with a Super PiggyBac transposase expression vector (System Biosciences) into HeLa TBK1 knockout cells and MC38 cells, respectively. Stable cell pools were generated by selection with blasticidin (10 µg/mL).

### CRISPR gene targeting

Guide RNAs were designed using the CRISPR design tool available at crispr.mit.edu. Single-guide RNAs (sgRNAs) targeting the human *TBK1* and *IKKε* genes were cloned into the pLC-GFP plasmid, which encodes Cas9 and GFP (kind gift from B. C. Bornhauser). Control cell lines were generated by transfection with the corresponding Cas9-expressing plasmid lacking sgRNA. Cells were transfected by electroporation, and 72 h post-transfection, GFP-positive cells were isolated by fluorescence-activated cell sorting (FACS). Single-cell clones were subsequently expanded and screened for gene knockout by immunoblotting for the respective proteins.

*hIKKε gRNA*: GTTGCGGGCCTTGTACACAC

*hTBK1 gRN*A: GCTACTGCAAATGTCTTTCG

### IRF3 and NF-κB reporter assay

HEK-Dual™ hTLR3 reporter cells (InvivoGen; Cat. #hkd-htlr3) were seeded in 96-well plates at a density of 1.0 × 10⁴ cells per well and allowed to adhere overnight. Cells were pre-treated with the TBK1 molecular glue CCT412020 for 12 h followed by stimulation with either poly(I:C) or TNF for an additional 6 h. After stimulation, 20 µL of culture supernatant was collected for reporter analysis. IRF3-dependent Lucia luciferase activity was measured using the Quanti-Luc™ 4 Lucia/Gaussia assay (InvivoGen; #rep-qlc4r5), and NF-κB-dependent SEAP activity was quantified using Quanti-Blue™ solution (InvivoGen; #rep-qbs3), according to the manufacturer’s instructions.

### Sulforhodamine B (SRB) clonogenic assay

Cells were seeded in either 6-well or 24-well plates at a density of 250 cells/well and left to adhere overnight. IFNβ was added for 12h pre-treatment, then TBK1 molecular glue for 6 h pre-treatment, before then remaining treatments were added simultaneously. Cells were incubated with treatments for 7 or 14 days with media and treatments being replenished every 3 days. Cells were fixed by adding cold 10% trichloroacetic acid per well for 1h at 4C (1 mL for 6-well plates, 500 µL for 24-well plates). Plates were then washed 4x in water and left to dry. SRB stain (SigmaAldrich; 0.057% w/v in 1% acetic acid) was added per well and incubated for 30 min at RT, then plates washed 4x in 1% acetic acid and left to dry. Plates were imaged using the Coomassie blue setting on a BioRad imager, and densitometry was calculated using ImageLab software (v6.1). Background intensity was subtracted from each value before normalising to DMSO control.

### Cell viability assays

Cell viability was measured by CellTiter-GLO assay (Promega) and was performed according to manufacturer’s instructions. Briefly, cells were seeded in 96-well plates and treated with required conditions, then cell survival was determined after the appropriate incubation time (24 or 48 h) by adding 20 µL CTG solution per well and measuring luminescence with a Victor X plate reader (PerkinElmer).

### Statistics & Reproducibility of *in vitro* cellular assays

Graphs and statistical analysis were performed using GraphPad Prism v9.5.1. The statistical analysis performed for each data set is described in the corresponding figure legend. Error bars indicate standard deviation (SD) or standard error of the mean (SEM), as indicated. All statistical tests were two-sided, and no statistical methods were used to predetermine sample size. No data were excluded from the analyses unless stated otherwise. Adjustment for multiple comparisons was performed unless indicated in the figure legends.

## Data availability

Source data will be made publicly available upon publication and will be deposited in Zenodo with a specific accession code.

## Acknowledgements

The Institute of Cancer Research Centre for Protein Degradation was established through support from philanthropists David and Ruth Hill.

Work in the Meier laboratory is funded by Breast Cancer Now as part of Programme Funding to the Breast Cancer Now Toby Robins Research Centre (CTR-QR14-007), CRUK programme funding (C26866/A24399), BBSRC (BB/W017261/1) and Worldwide Cancer Research (23-0146). We acknowledge NHS funding to the NIHR Biomedical Research Centre. This study represents independent research supported by the National Institute for Health Research (NIHR) Biomedical Research Centre at The Royal Marsden NHS Foundation Trust and The Institute of Cancer Research, London. The views expressed are those of the authors and not necessarily those of the NIHR or the Department of Health and Social Care.

NEOsphere Biotechnologies GmbH gratefully acknowledges funding for this research from the German Federal Ministry of Education and Research (BMBF, grant number 16LW0372). We thank Dr. Jutta Fritz for her support in project management, and Ines Scheller for providing tools for proteomics data analysis. We further thank Denis Bartoschek, Anastasia H. Bednarz, Sophie Machata, and Tobias Graef for performing proteomics and global ubiquitinomics experiments and providing technical support.

## Author contributions

H.D. and Z.R. conceived the study, designed the experiments, analysed and interpreted the data, and wrote the manuscript.

P.R.A.Z. designed the proteomics experiments, analysed and interpreted the data, prepared **Fig. 2 and Suppl. Fig. 1**, and wrote the proteomics section of the manuscript. M.S. co-developed the AE-MS assay, analysed and interpreted the proteomics data. D.W. co-developed the AE-MS assay and performed the experiments. U.O. analysed and interpreted the proteomics data. B.Sch. and B.Sh. developed the proteomics data analysis pipeline, analysed and interpreted proteomics data. H.D. conceived the study, designed the experiments, analysed and interpreted the data, and was involved in manuscript writing and editing. S.J. performed and analysed the mutants. P.M., T.T., R.W. and S.J. designed and performed experiments in **Fig. 3D, E, I** and K. P.M., T.T., R.W., and I.F. designed, performed and analysed experiments in Figure 5 and Suppl. Fig. 5. S.K performed the *in vitro* hit validation and mechanism of action studies in Figure 5 and Suppl. Fig. 2. S.K and A.K designed the experiments, analysed and interpreted data, prepared figures and were involved in manuscript writing. P.C.M.A and S.T.H. cloned, produced and purified recombinant proteins. S.T.H. performed SEC ternary complex formation experiments, prepared cryo-EM grids, collected cryo-EM data and solved cryo-EM structures. S.T.H. and Y.-V.L.B. analysed and interpreted the data, prepared the figures, and were involved in manuscript writing and editing. R.L.M.v.M. designed and led structural biology experiments and was involved in manuscript writing. N.S.A. developed cheminformatics workflows for the molecular glue library design and selection, prepared figures and was involved in manuscript writing. A.S. directed the cheminformatics design strategy. J.J.C. designed and selected the molecular glue library, prepared figures and was involved in manuscript writing. R.J.H.W was involved in manuscript writing. K.M and M.T.W designed and implemented the G-loop identification code, and analysed and interpreted the data. M.T.W was involved in figure preparation and manuscript writing. F.D managed the project and was involved in manuscript editing.

All authors read and approved the final manuscript

## Supplementary Figures

**Suppl. Fig. 1.**
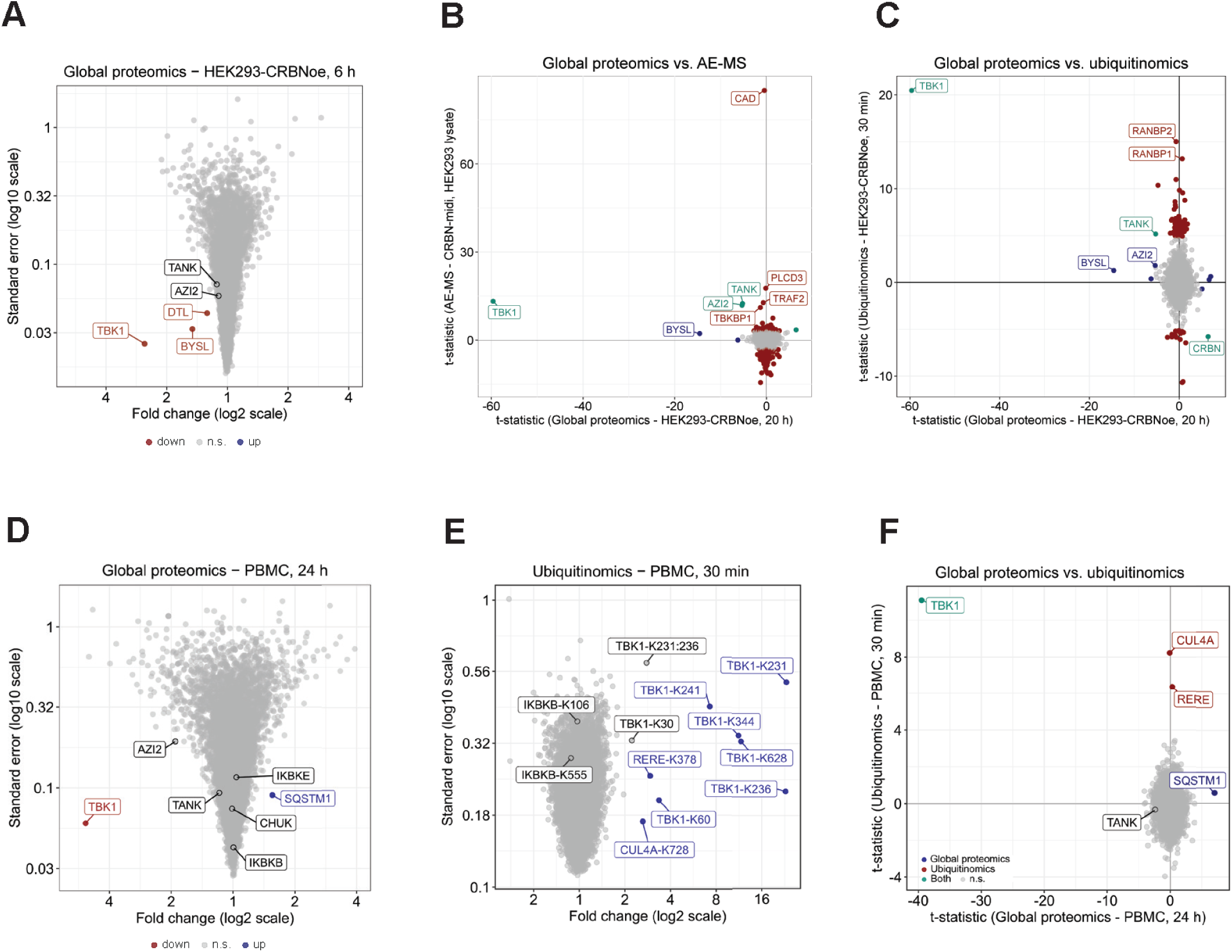
Identification and characterization of a TBK1 molecular glue degrader using high-throughput proteomics. In all volcano plots (A, D and F), fold-changes relative to vehicle (x-axis, log_2_ scale) and standard errors (y-axis, log_10_ scale) are shown. Up- and downregulated proteins (q < 0.01) or ubiquitination sites (q < 0.05) are coloured in blue and red, respectively. Not significant (n.s.) regulations are coloured grey. In comparison plots (B, C and E), t-statistics for regulations in global proteomics data (x-axis)) and regulations in a second assay type (AE-MS or ubiquitinomics, y-axis) are plotted against each other. For this purpose, ubiquitination sites mapped to the same protein were averaged, and only significantly upregulated sites were included in the analysis (C and E). Statistically significant up- and downregulations in the proteome are coloured in blue (q < 0.01), whereas those in the ubiquitinomics (q < 0.05) or AE-MS interactomics data (q < 0.01) are labelled in red. Proteins significantly down-regulated in the proteome and significantly up-regulated in either the AE-MS interactomics or ubiquitinomics data are highlighted in turquoise, not significant regulations are coloured grey. **(A)** Global proteomic profile of CCT412020 in HEK293-CRBNoe cells (10 µM, 6 h) shows weaker TBK1 degradation compared to the 20 h timepoint (log*_2_*FC = -2.18). **(B)** t-Statistical comparison of global proteomics data from CCT412020-treated HEK293-CRBNoe cells (10 µM, 20 h) and AE-MS data of CCT412020-induced (10 µM) protein binding to immobilised CRBN^midi^ in HEK293 lysate. **(C)** t-Statistical comparison of global proteomics and ubiquitinomics data from HEK293-CRBNoe cells treated with 10 µM CCT412020 for 20 h and 30 min, respectively. **(D)** Global proteomics profile of CCT412020 in PBMCs (10 µM, 24 h). **(E)** Global ubiquitinomics by K-GG remnant peptide profiling reveals pronounced ubiquitination of multiple TBK1 lysine residues induced by CCT412020 (10 µM, 30 min) in PBMCs. **(F)** t-statistical comparison of global proteomics (10 µM, 24 h) and ubiquitinomics data (10 µM, 30 min) from PBMCs treated with 10 µM CCT412020 for 20 h and 30 min, respectively.

**Suppl. Fig. 2.**
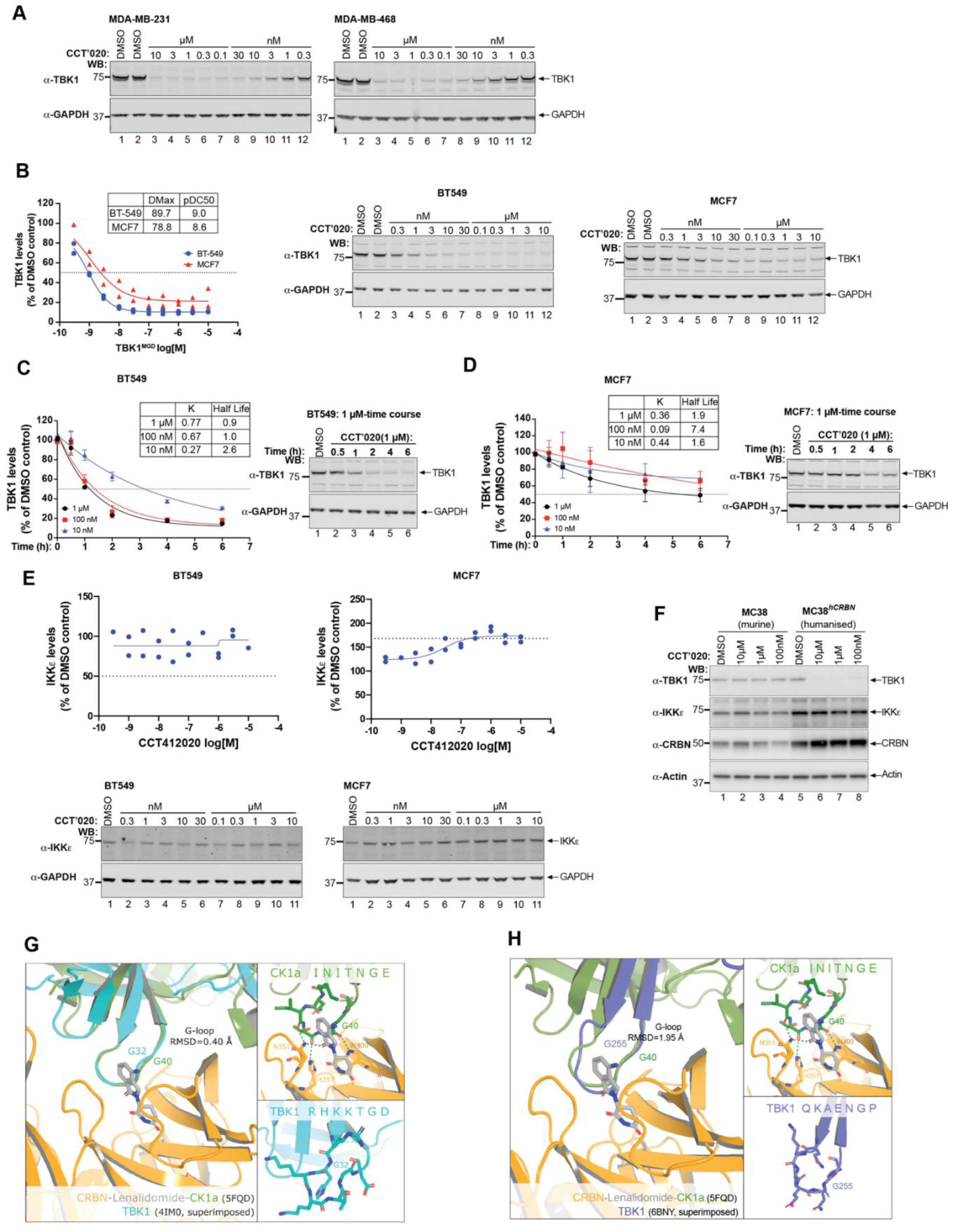
Pharmacological profiling of CCT412020 in selected breast cancer cell lines. (**A**) Representative degradation western blots in in breast cancer cell lines MDA-MB-231 and MDA-MB-468 for Fig. 3F. (**B**) CCT412020 has a strong TBK1 degradation potency and D_max_ in breast cancer cell lines, BT549 (pDC_50_Abs 9.0, D_max_ 90%), MCF7 (pDC_50_Abs 8.6, D_max_ 79%). (**C**) TBK1 time-course degradation study in BT549 cells demonstrating fast degradation kinetics resulting in over 70% degradation within 2 h. (**D**) Degradation time-course in MCF7 cells demonstrating 50% degradation within 6 h, highlighting differences between cell lines. (**E**) CCT412020 does not degrade IKKe in BT549 and MCF7 cells, dose-response study. Small increase of IKKe is observed in MCF7 cells. (**F**) Western blot analysis of MC38 parental and MC38^hCRBN^ cells treated with CCT412020 (18h). (**G**) Predicted G-loop degron from TBK1 (cyan) aligned to the CK1α degron (green) in complex with CRBN (orange) bound to lenalidomide (grey). Top inset: CK1α degron sequence and structure highlighting hydrogen-bond interactions between the degron, CRBN, and lenalidomide (PDB 5FQD). Bottom inset: predicted TBK1 degron sequence and structure from the TBK1 crystal structure (PDB 4IM0). (**H**) Alternative Predicted G-loop degron from TBK1 (purple) aligned to the CK1α degron (green) in complex with CRBN (orange) bound to lenalidomide (grey). Top inset: CK1α degron sequence and structure highlighting hydrogen-bond interactions between the degron, CRBN, and lenalidomide (PDB 5FQD). Bottom inset: predicted TBK1 degron sequence and structure from the TBK1 crystal structure (PDB 6BNY).

**Suppl. Fig. 3.**
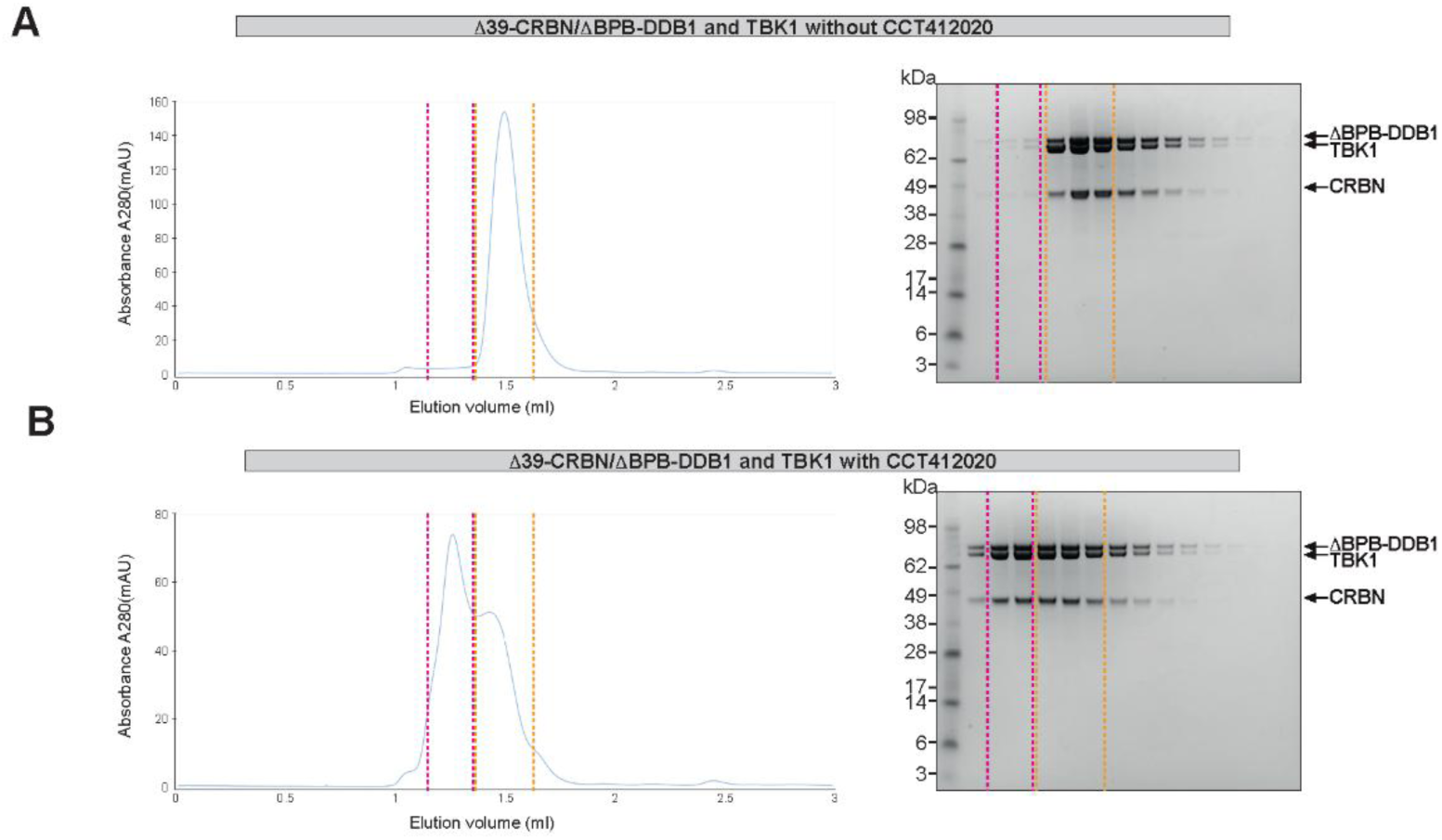
Size exclusion chromatography (SEC) analysis of complex formation between CRBN/DDB1 and TBK1 induced by molecular glue CCT412020. **(A)** Left-Analytical SEC chromatogram of Δ39-CRBN/ΔBPB-DDB1 and TBK1 in the absence of CCT412020. Right-SDS-PAGE analysis of the resulting fractions. **(B)** Left-Analytical SEC chromatogram of Δ39-CRBN/ΔBPB-DDB1 and TBK1 in the presence of CCT412020. Right-SDS-PAGE analysis of the resulting fractions. Dashed purple and orange lines highlight the equivalent regions of the SEC chromatograms and the SDS-PAGEs. In the absence of CCT412020, only one main peak is seen in the chromatogram (highlighted by the orange dashed lines), which contains Δ39-CRBN, ΔBPB-DDB1 and TBK1, as the Δ39-CRBN/ΔBPB-DDB1 complex and the TBK1 dimer have roughly the same molecular weight (141 and 152 kDa, respectively). In the presence of CCT412020, a new peak is observed in the chromatogram at lower retention volume (highlighted by the purple dashed lines), which still contains Δ39-CRBN, ΔBPB-DDB1 and TBK1, demonstrating the formation of a complex between Δ39-CRBN/ΔBPB-DDB1 and TBK1 in the presence of CCT412020.

**Suppl. Fig. 4.**
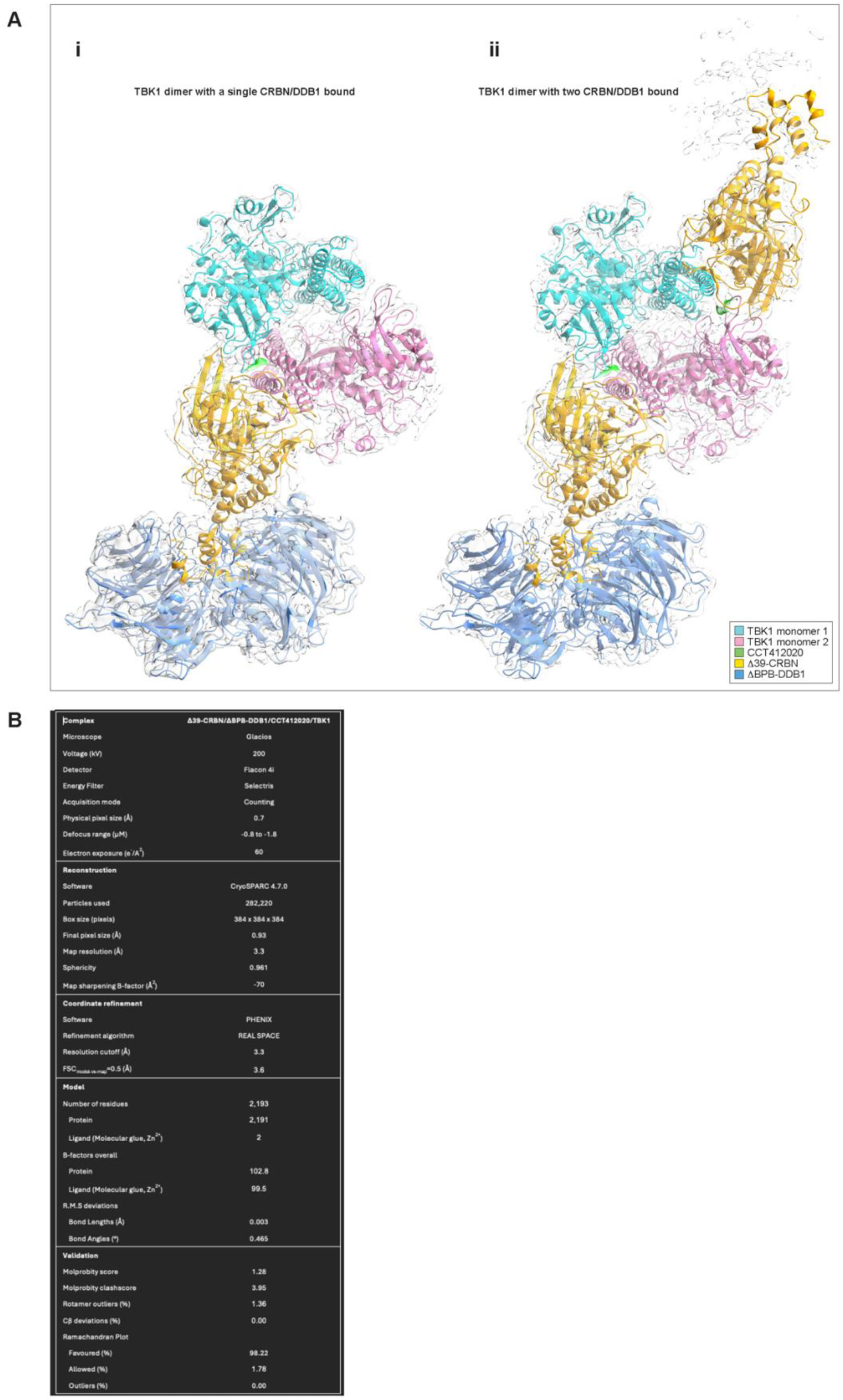
Two different stoichiometries of CRBN/DDB1 bound to the TBK1 dimer were observed during cryo-EM analysis. **(A)** Comparison of two models of the Δ39-CRBN/ΔBPB-DDB1/CCT412020/TBK1 complex obtained by cryo-EM. i, The TBK1 dimer bound to a single CRBN/DDB1 complex. ii, The TBK1 dimer bound to two CRBN/DDB1 complexes on opposite sides of the TBK1 dimer. The density for the second DDB1 component (top right) was insufficient to allow reliable model docking due to poor alignment in this region. The TBK1 dimer (cyan and pink), CRBN (yellow) and DDB1 (blue) are represented as cartoon traces. The EM maps are represented as a semi-transparent grey surfaces. Structural figures were generated using ChimeraX. **(B)** Cryo-EM data collection, 3D reconstruction, refinement and validation statistics.

**Suppl. Fig. 5.**
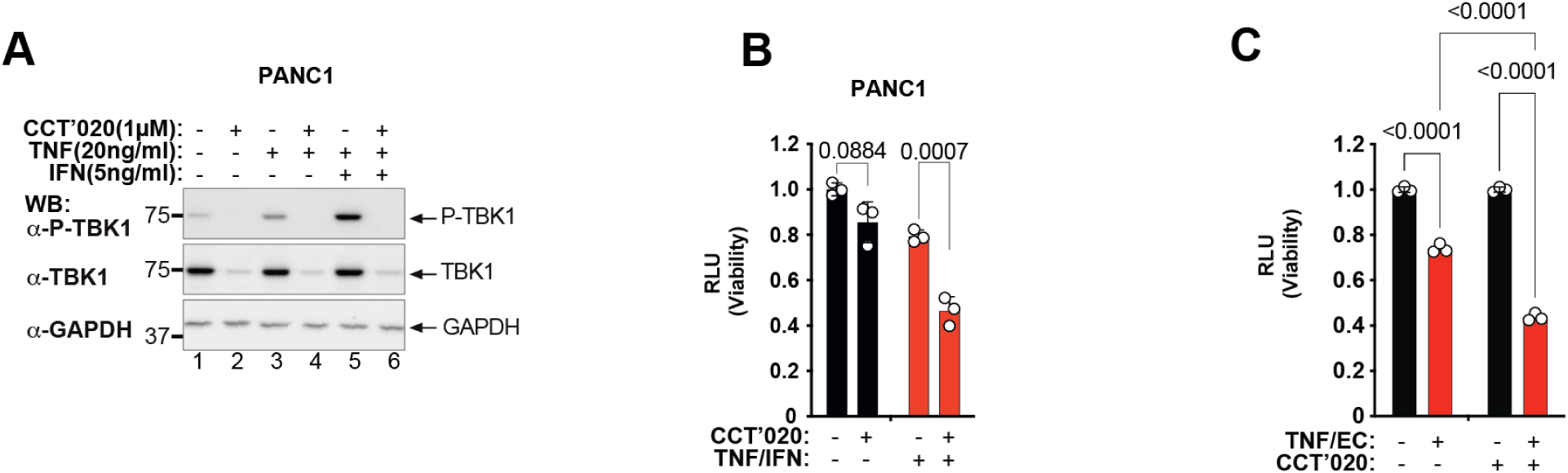
TNF and IFN sensitisation of cancer cell line treated with the TBK1 molecular glue CCT412020. **(A)** Western blot analysis of total TBK1 and phosphorylated TBK1 (p-TBK1) in PANC-1 cells treated with the TBK1 molecular glue CCT412020 (1 µM) in the presence or absence of TNF and/or IFNγ. **(B)** Cell viability assay in PANC-1 cells treated with the TBK1 molecular glue CCT412020 (1 µM) in the presence or absence of TNF (10 ng/ml)/IFNγ (5 ng/ml). Cells were treated for 48h, and viability was quantified following treatment. **(C)** Cell viability assay in MC38^hCRBN^ cells treated with TNF and the pan-caspase inhibitor emricasan to induce necroptosis, in the presence of either the TBK1 molecular glue CCT412020 (1 µM). Cell viability was quantified following treatment as indicated.

